# Enhanced CICR activity reduces ER Ca^2+^ level in cells expressing CPVT-linked mutant RyR2

**DOI:** 10.1101/2021.01.16.426980

**Authors:** Nagomi Kurebayashi, Takashi Murayama, Ryosaku Ohta, Junji Suzuki, Kazunori Kanemaru, Seiko Ohno, Minoru Horie, Masamitsu Iino, Fumiyoshi Yamashita, Takashi Sakurai

**Affiliations:** Department of Cellular and Molecular Pharmacology, Juntendo University Graduate School of Medicine, Tokyo, Japan; Department of Drug Delivery Research, Graduate School of Pharmaceutical Sciences, Kyoto University, Kyoto, Japan; Department of Physiology, University of California San Francisco, San Francisco, CA, USA; Division of Cellular and Molecular Pharmacology, Nihon University School of Medicine, Tokyo, Japan; Department of Bioscience and Genetics, National Cerebral and Cardiovascular Center Research Institute, Osaka, Japan; Center for Epidemiologic Research in Asia, Shiga University of Medical Science, Otsu, Shiga, Japan

**Author notes:** These authors contributed equally to this work (Nagomi Kurebayashi, Takashi Murayama, Ryosaku Ohta, Fumiyoshi Yamashita). **Corresponding authors:** Nagomi Kurebayashi, Ph.D., Department of Cellular and Molecular Pharmacology, Juntendo University Graduate School of Medicine, Tokyo 113-8421, Japan, Fumiyoshi Yamashita, Ph.D., Department of Drug Delivery Research, Graduate School of Pharmaceutical Sciences, Kyoto University, Kyoto 606-8501, Japan.

## Abstract

Type 2 ryanodine receptor (RyR2) is a cardiac Ca^2+^ release channel in the endoplasmic reticulum (ER). Mutations in RyR2 are linked to catecholaminergic polymorphic ventricular tachycardia (CPVT), which is considered to be associated with enhanced spontaneous Ca^2+^ release. This spontaneous Ca^2+^ release tends to occur when ER Ca^2+^ ([Ca^2+^]_ER_) reaches a certain threshold level, and CPVT mutations are reported to lower this threshold. There are two explanations for this lowered threshold: the mutations increase sensitivity to luminal Ca^2+^ or they enhance cytosolic [Ca^2+^] ([Ca^2+^]_cyt_)-induced Ca^2+^ release (CICR) activity. However, no quantitative analysis of this issue has been performed so far. Here, we quantitatively explored how the change in CICR activity of RyR2 affects the threshold [Ca^2+^]_ER_ experimentally and by model-based simulation. Wild-type (WT) and CPVT-linked mutant RyR2s were expressed in HEK293 cells. [Ca^2+^]_cyt_ and [Ca^2+^]_ER_ measurements with Ca^2+^ indicators revealed that CPVT RyR2 cells showed higher oscillation frequency and lower threshold [Ca^2+^]_ER_ in a mutation-specific manner compared with WT cells. The CICR activity of mutant RyR2s was assessed by Ca^2+^-dependent [^3^H]ryanodine binding and parameter analysis. CICR activity at resting [Ca^2+^]_cyt_, A_7.0_, was higher in CPVT mutants than in WT and a strong inverse correlation was found between threshold [Ca^2+^]_ER_ and A_7.0_. Interestingly, lowering RyR2 expression increased threshold [Ca^2+^]_ER_, suggesting that the threshold [Ca^2+^]_ER_ depends on net Ca^2+^ release rate via RyR2, a product of A_7.0_ for each mutant and the density of RyR2 molecules. A model-based simulation successfully reproduced the [Ca^2+^]_cyt_ and [Ca^2+^]_ER_ changes. Interestingly, the CICR activity associated with specific mutations correlated well with the age of onset of the disease in CPVT patients carrying the mutations. Our data suggest that the reduction in threshold [Ca^2+^]_ER_ for spontaneous Ca^2+^ release by CPVT mutation is explained by enhanced CICR activity without considering a change in the [Ca^2+^]_ER_ sensitivity of RyR2.

**Summary:** CPVT-linked RyR2 mutations are prone to induce spontaneous Ca^2+^ release from ER, which is strongly associated with arrhythmias. Kurebayashi et al. quantitatively explore how the changes in CICR activity by RyR2 mutations affect spontaneous Ca^2+^ experimentally and by model simulation.

## Introduction

The type 2 ryanodine receptor (RyR2) is a Ca^2+^ release channel in the sarcoplasmic/endoplasmic reticulum (SR/ER) that plays an indispensable role in excitation–contraction coupling in the heart. In cardiac myocytes, RyR2 is activated by Ca^2+^ influx through L-type Ca^2+^ channels during action potential with a cytosolic Ca^2+^ ([Ca^2+^]_cyt_)-induced Ca^2+^ release (CICR) mechanism to release more Ca^2+^, which in turn causes muscle contraction (Fabiato, 1983; Bers, 2002). In contrast, spontaneous Ca^2+^ release such as Ca^2+^ waves, which lead to delayed afterdepolarization (DAD) of the plasma membrane via a Na^+^–Ca^2+^ exchange reaction, often results in triggered activity (Tsien et al., 1979; Lakatta, 1992). The spontaneous Ca^2+^ release can be seen when cardiomyocytes are Ca^2+^-overloaded, even in healthy hearts, but is more likely to occur in myocardium in heart failure patients and carriers of RyR2 mutations associated with sudden cardiac death syndrome.

Mutations in RyR2 have been linked to several types of arrhythmogenic disease, such as catecholaminergic polymorphic ventricular tachycardia (CPVT), left ventricular non-compaction (LVNC), and idiopathic ventricular fibrillation (IVF) (Priori et al., 2001; Tester et al., 2004; Medeiros-Domingo et al., 2009; Priori and Chen, 2011; Kawamura et al., 2013; Fujii et al., 2017; Uehara et al., 2017; Nozaki et al., 2020), and nearly 300 arrhythmogenic mutations have been reported to date. CPVT is the most common RyR2-related arrhythmogenic disorder and is induced in response to sympathetic nerve activation without structural abnormality of the heart (Priori et al., 2001). CPVT-linked RyR2 mutations characterized so far were associated with gain-of-function phenotypes, which are prone to induce spontaneous Ca^2+^ release from SR.

Two Ca^2+^-dependent regulatory mechanisms of RyR2—the regulation from the luminal side of the ER and that from the cytoplasmic side—may be involved in the spontaneous Ca^2+^ release. Many reports have described that luminal [Ca^2+^] plays a role in controlling SR Ca^2+^ release (Bassani et al., 1995; Lukyanenko et al., 1996; Sitsapesan and Williams, 1997). Single-channel analyses have indicated that ER luminal Ca^2+^ activates RyR2 at around 10^−3^ M in the presence of calsequestrin 2 (CSQ2) and at 10^−2^ M in its absence (Qin et al., 2008). In contrast, the role of cytosolic Ca^2+^ is known as the CICR mechanism (Endo, 1977; Fabiato, 1983; Murayama and Kurebayashi, 2011; Guo et al., 2012; Rios, 2018). The cytoplasmic Ca^2+^ activates RyR2 at low concentrations (<~10^−4^ M) of Ca^2+^ and suppresses it at high concentrations (>~10^−3^M), and consequently RyR2 channel activity displays a bell-shaped Ca^2+^ dependence on cytoplasmic Ca^2+^ (Murayama and Kurebayashi, 2011; Rios, 2018). Therefore, a small local Ca^2+^ leak from the ER, which is more likely to occur at high [Ca^2+^]_ER_, may activate RyR2 from the cytoplasmic side to trigger massive Ca^2+^ release by the positive feedback nature of CICR.

At present, there is controversy over whether CPVT mutations primarily affect the cytoplasmic or luminal regulation of RyR2. Chen and colleagues indicated that the spontaneous Ca^2+^ release, called store-overload-induced Ca^2+^ release (SOICR), in RyR2-expressing HEK293 cells occurs when [Ca^2+^]_ER_ reaches a certain critical threshold level (Jiang et al., 2004; Jiang et al., 2005) and found that the CPVT-linked RyR2 cells showed a lowered threshold [Ca^2+^]_ER_ for spontaneous Ca^2+^ release compared with WT RyR2 (Jones et al., 2008). They reported that some of the CPVT-linked RyR2 channels (e.g., R2474S and R4497C) showed increased luminal Ca^2+^ sensitivity with no significant effects on cytoplasmic Ca^2+^ dependence (Jiang et al., 2005), suggesting that the lowered threshold [Ca^2+^]_ER_ with these CPVT mutations is due to sensitization to luminal Ca^2+^ but not to cytosolic Ca^2+^. In contrast, Wehrens et al. reported that the same mutation, R2474S, showed increased cytoplasmic Ca^2+^ sensitivity via the phosphorylation of RyR2 by protein kinase A due to the dissociation of FKBP12.6 (Wehrens et al., 2003). We also found that R2474S showed higher CICR activity at resting [Ca^2+^]_cyt_ (Uehara et al., 2017). The reason for this discrepancy may be better understood by a more quantitative analysis․.

The HEK293 cell expression system allows both functional and biochemical analyses of exogenously expressed RyRs (Jiang et al., 2002; Jiang et al., 2004; Gomez and Yamaguchi, 2014; Murayama et al., 2015). In HEK293 cells, regulatory proteins specific to myocardium, such as calsequestrin and FKBP12.6, are absent, so the use of these cells enables evaluation of the direct effect of mutations on RyR2 activity. We quantitatively evaluated the Ca^2+^ release activity of RyR1 using the HEK293 expression system and showed that [Ca^2+^]_ER_ signals are well correlated with the [Ca^2+^]_cyt_-dependent [^3^H]ryanodine binding activity of RyR1 (Murayama et al., 2015; Murayama et al., 2016). In this study, we investigated the correlation between CICR activity and [Ca^2+^]_ER_ in HEK293 cells using 10 CPVT and 6 artificial RyR2 mutations. We further validated this correlation using a mathematical model. Our results suggest that the changes in threshold [Ca^2+^]_ER_ for spontaneous Ca^2+^ release by CPVT mutation are explained by CICR activity without considering a change in the [Ca^2+^]_ER_ sensitivity of RyR2.

## Methods

### Generation of stable inducible HEK293 cell lines

Full-length mouse RyR2 cDNA was constructed from cDNA fragments that were PCR-amplified from mouse ventricle and then cloned into a tetracycline-induced expression vector (pcDNA5/FRT/TO; Life Technologies, CA, USA) (Fujii et al., 2017; Uehara et al., 2017). Mutations corresponding to V2321M, R2474S, D3638A, Q4201R, K4932R, R4497C, K4751Q, H4762P, K4805R, and I4867M were introduced by inverse PCR and confirmed by DNA sequencing (Table 1). The expression vector was co-transfected with pOG44 into Flp-In T-REx HEK293 cells (Life Technologies), in accordance with the manufacturer’s instructions. Clones with suitable doxycycline-induced expression of RyR2 were selected and used for the experiments.

**Table 1.**
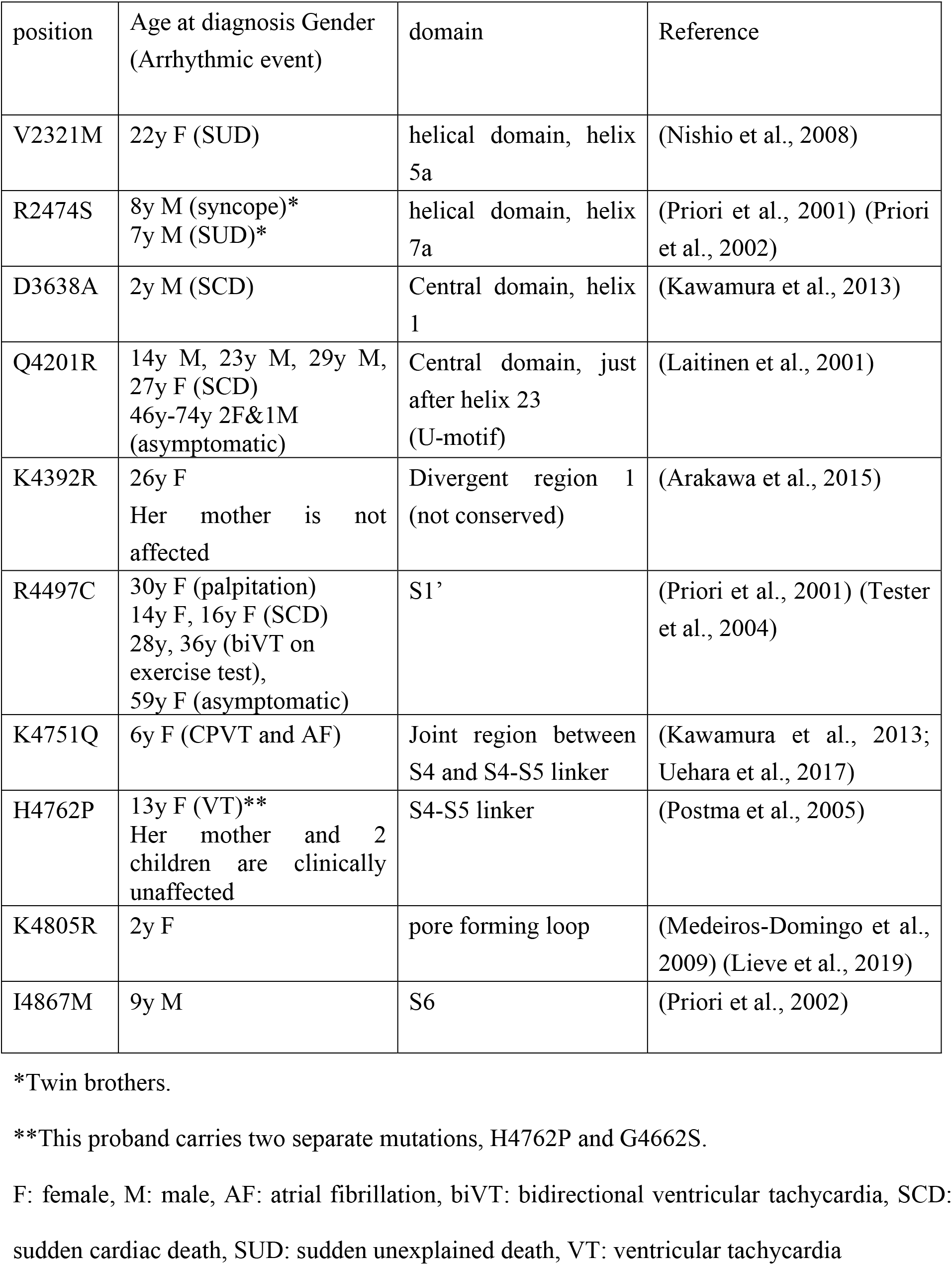
List of CPVT mutations examined in this study.

| position | Age at diagnosis Gender<br>(Arrhythmic event) | domain | Reference |
| --- | --- | --- | --- |
| V2321M | 22y F (SUD) | helical domain, helix 5a | (Nishio et al., 2008) |
| R2474S | 8y M (syncope)*<br>7y M (SUD)* | helical domain, helix 7a | (Priori et al., 2001) (Priori et al., 2002) |
| D3638A | 2y M (SCD) | Central domain, helix 1 | (Kawamura et al., 2013) |
| Q4201R | 14y M, 23y M, 29y M, 27y F (SCD)<br>46y-74y 2F&1M (asymptomatic) | Central domain, just after helix 23 (U-motif) | (Laitinen et al., 2001) |
| K4392R | 26y F<br>Her mother is not affected | Divergent region 1 (not conserved) | (Arakawa et al., 2015) |
| R4497C | 30y F (palpitation)<br>14y F, 16y F (SCD)<br>28y, 36y (biVT on exercise test),<br>59y F (asymptomatic) | S1' | (Priori et al., 2001) (Tester et al., 2004) |
| K4751Q | 6y F (CPVT and AF) | Joint region between S4 and S4-S5 linker | (Kawamura et al., 2013; Uehara et al., 2017) |
| H4762P | 13y F (VT)**<br>Her mother and 2 children are clinically unaffected | S4-S5 linker | (Postma et al., 2005) |
| K4805R | 2y F | pore forming loop | (Medeiros-Domingo et al., 2009) (Lieve et al., 2019) |
| I4867M | 9y M | S6 | (Priori et al., 2002) |

\*Twin brothers.
\*\*This proband carries two separate mutations, H4762P and G4662S.
F: female, M: male, AF: atrial fibrillation, biVT: bidirectional ventricular tachycardia, SCD: sudden cardiac death, SUD: sudden unexplained death, VT: ventricular tachycardia

### Single-cell Ca^2+^ imaging

HEK293 cells grown on a glass-bottomed dish were treated with doxycycline 26–28 h prior to measurement to induce the expression of RyR2, unless otherwise noted. Single-cell Ca^2+^ imaging was carried out in HEK293 cells expressing WT or mutant RyR2 in HEPES-buffered Krebs solution (140 mM NaCl, 5 mM KCl, 2 mM CaCl_2_, 1 mM MgCl_2_, 11 mM glucose, 5 mM HEPES, pH 7.4) (Murayama et al., 2015; Uehara et al., 2017). For cytosolic Ca^2+^ measurements, cells were loaded with 4 μM fluo-4 AM in culture medium for 30 min at 37°C and then incubated with Krebs solution. Fluo-4 was excited at 488 nm through a 20× objective lens and emitting light at 525 nm was captured with an EM-CCD camera at 700 ms intervals (Model 8509; Hamamatsu Photonics, Hamamatsu, Japan). The fluorescence signal (*F*) of fluo-4 in normal Krebs solution in individual cells was determined using region of interest (ROI) analysis, was subtracted cell-free background fluorescence, and was normalized to the maximal fluorescence intensity (*F*_max_) obtained with a 20Ca-Krebs solution (140 mM NaCl, 5 mM KCl, 20 mM CaCl_2_, 1 mM MgCl_2_, 11 mM glucose, 20 μM ionomycin, 5 mM HEPES, pH 7.4) at the end of each experiment. All measurements were carried out at 26 °C by perfusing solutions through an in-line solution heater/cooler (Warner Instruments, Holliston, MA, USA). [Ca^2+^]_cyt_ was calculated using the parameters of effective *K*_D_ = 2.2 μM and *n* = 1 *in situ* (Harkins et al., 1993; Nelson et al., 2014).

[Ca^2+^]_cyt_ and [Ca^2+^]_ER_ were simultaneously monitored using genetically encoded Ca^2+^ indicators, G-GECO1.1 (Zhao et al., 2011) and R-CEPIA1er, (Suzuki et al., 2014; Murayama et al., 2015; Uehara et al., 2017), respectively. Cells were transfected with G-GECO1.1 and R-CEPIA1er cDNA 26–28 h before measurements. Doxycycline was added to the medium at the same time as transfection in most experiments, and after transfection in the experiments for Figure 7. Cytosolic and ER Ca^2+^ signals were obtained in normal Krebs solution for 5 min and then in 10 mM caffeine containing Krebs solution. At the end of the measurement, the cells were perfused with the following solutions: 0Ca-Krebs solution (140 mM NaCl, 5 mM KCl, 1 mM MgCl_2_, 11 mM glucose, 5 mM HEPES, pH 7.4), BAPTA-0Ca-Krebs solution [140 mM NaCl, 5 mM KCl, 5 mM 1,2-bis(o-aminophenoxy)ethane-N,N,N′,N′-tetraacetic acid (BAPTA), 1 mM MgCl_2_, 11 mM glucose, 20 μM ionomycin, 20 μM cyclopiazonic acid, 5 mM HEPES, pH 7.4], 0Ca-Krebs solution, and then 20Ca-Krebs solution. *Fmin* and *F*_max_ values were obtained with the BAPTA-0Ca-Krebs solution and 20Ca-Krebs solution, respectively. Because G-GECO1.1 signals have a high Hill coefficient (*n* = 3.38) (Suzuki et al., 2014), which makes the calculation of [Ca^2+^]_cyt_ difficult, only the R-CEPIA1er signal was used for the calculation of [Ca^2+^]_ER_. [Ca^2+^]_ER_ was calculated using parameters obtained by *in situ* titration (*K*_D_ = 565 μM, *n* = 1.7) (Suzuki et al., 2014).

### [^3^H]Ryanodine binding and parameter analysis

Ca^2+^-dependent [^3^H]ryanodine binding was performed as described previously (Fujii et al., 2017; Nozaki et al., 2020). Briefly, microsomes isolated from HEK293 cells were incubated for 1 h at 25 °C with 5 nM [^3^H]ryanodine in a medium containing 0.17 M NaCl, 20 mM 3-(*N*-morpholino)-2-hydroxypropanesulfonic acid (MOPSO), pH 7.0, 2 mM dithiothreitol, 1 mM AMP, 1 mM MgCl_2_, and various concentrations of free Ca^2+^ buffered with 10 mM ethylene glycol-bis(2-aminoethylether)-N,N,N′,N′-tetraacetic acid (EGTA). Free Ca^2+^ concentrations were calculated using WEBMAXC STANDARD (https://somapp.ucdmc.ucdavis.edu/pharmacology/bers/maxchelator/webmaxc/webmaxcS.htm) (Bers et al., 2010). The protein-bound [^3^H]ryanodine was separated by filtering through polyethyleneimine-treated GF/B filters using Micro 96 Cell Harvester (Skatron Instruments, Lier, Norway). Nonspecific binding was determined in the presence of 20 μM unlabeled ryanodine. The [^3^H]ryanodine binding data (*B*) were normalized to the maximum number of functional channels (*B*_max_), which was separately determined by Scatchard plot analysis using various concentrations (3–20 nM) of [^3^H]ryanodine in a high-salt medium containing 1 M NaCl. The resultant *B*/*B*_max_ represents the averaged activity of each mutant.

The Ca^2+^-dependent [^3^H]ryanodine binding data were parameterized with the following model, in which the channel activity is determined by three independent parameters (*A*_max_, *K*_ACa_, and *K*_ICa_):

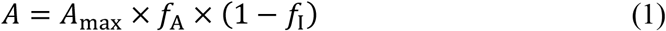

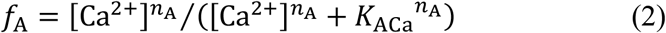

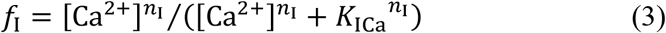

where *A* is the activity at the specified Ca^2+^, *A*_max_ is the gain that determines the maximal attainable activity, and *f*_A_ and *f*_I_ are fractions of the activating Ca^2+^ site (A-site) and inactivating Ca^2+^ site (I-site) occupied by Ca^2+^, respectively. *KACa* and *KICa* are dissociation constants, and *n*_A_ and *n*_I_ are Hill coefficients for Ca^2+^ of A- and I-sites, respectively. We set the Hill coefficients (*n*_A_ = 2.0 and *n*_I_ = 1.0) to values that maximize the sum of *R*^2^ values for curve fitting (Murayama and Kurebayashi, 2011; Fujii et al., 2017). Curve fitting was performed using Prism 8 software (GraphPad Software, San Diego, CA, USA).

For estimation of ryanodine binding at resting Ca^2+^, *A* at pCa 7.0 (A_7.0_) for each mutant was calculated by Equations (1)–(3) using the obtained parameters (*KACa*, *KICa*, and *A*_max_) (Table 2), in which free [Ca^2+^] was set at 100 nM.

**Table 2.** Parameters for CICR activities of CPVT and artificial mutants.

| | $A_{max}$ | $K_{Aca}$ (mM) | $K_{ICa}$ (mM) |
| --- | --- | --- | --- |
| Mock | 0 |  |  |
| WT | 0.1248 | 0.0162 | 3.002 |
| V2321M | 0.1847 | 1.10E-02 | 4.847 |
| R2474S | 0.2175 | 7.03E-03 | 11.31 |
| D3638A | 0.1214 | 1.10E-02 | 2.058 |
| Q4201R | 0.1473 | 1.04E-02 | 4.454 |
| K4392R | 0.1324 | 0.0180 | 2.759 |
| R4497C | 0.1779 | 1.29E-02 | 5.553 |
| K4751Q | 0.2733 | 7.65E-03 | 1000 |
| H4762P | 0.09848 | 2.94E-03 | 19.5 |
| K4805R | 0.2857 | 7.35E-03 | 159.5 |
| I4867M | 0.08865 | 8.96E-03 | 1.832 |
| T4754A | 0.1245 | 0.0203 | 2.178 |
| I4755A | 0.06616 | 8.18E-03 | 1.829 |
| L4756A | 0.1925 | 2.22E-03 | 741.6 |
| S4757A | 0.1684 | 1.38E-02 | 6.967 |
| S4758A | 0.05471 | 0.01953 | 3.245 |
| T4754I | 0.2953 | 8.40E-03 | 17.13 |

### Western blotting

Microsomal proteins were separated by sodium dodecyl sulfate polyacrylamide gel electrophoresis and transferred onto a polyvinylidene fluoride membrane. Western blotting was performed using antibodies for RyR2 (Chugun et al., 2007) and calnexin (C4731; Sigma-Aldrich, St. Louis, MO, USA).

### Mathematical model simulation

To mathematically describe calcium dynamics, cells were assumed to comprise the cytosolic and ER compartments (see Figure 8A), where the transport of Ca^2+^ between these compartments involves Ca^2+^-induced Ca^2+^ release (CICR), saturable ER Ca^2+^ uptake, and passive leakage. We assumed herewith that intra-compartmental Ca^2+^ reaches an equilibrium instantly. Cellular Ca^2+^ influx is assumed to obey zero-order kinetics provided that the extracellular bulk concentration is constant. In contrast, extracellular Ca^2+^ efflux is assumed to obey Michaelis–Menten kinetics depending on the cytoplasmic Ca^2+^ concentration. Additionally, the presence of Ca^2+^-buffering proteins was considered. Taking this together with the assumption that CICR activity is proportional to in vitro ryanodine binding, the calcium dynamics model gives the following mass balance equations:

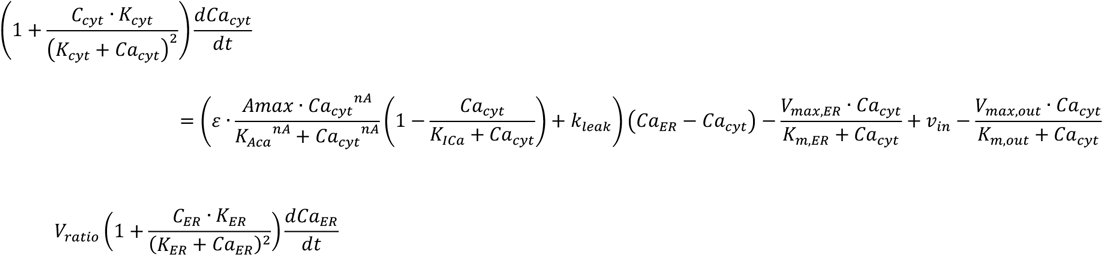

where *Cacyt* and *CaER* are the free Ca^2+^ concentrations in cytosol and ER *Vratio* is the ratio of the ER to cytosol volumes; *Ccyt* and *Kcyt* are the total concentration of cytosolic Ca^2+^ buffering site and its dissociation constant; *CER* and *KER* are the total concentration and the dissociation constant for calreticulin in the ER (Means et al., 2006); *Amax*, *KACa*, *KICa*, and *nA* are the maximum binding, the dissociation constant for activating Ca^2+^, the dissociation constant for inhibiting Ca^2+^, and the Hill coefficient for ryanodine binding; *∊* is a correction factor between *Amax* and maximal activity for CICR; *kleak* is the permeability constant for *Ca^2+^* leakage across ER membranes; *Vmax,ER* and *Km,ER* are the maximal rate and Michaelis–Menten constant for ER uptake; *vin* is the zero-order influx rate from extracellular fluid; and *Vmax,out* and *Km,out* are the maximal rate and Michaelis–Menten constant for extracellular efflux. The *Vratio* value used for the simulation was 0.03, which is close to that in cardiomyocytes (Shannon et al., 2004). *Ccyt* and *Kcyt* were 11 mM and 0.3 mM respectively, by simplifying multiple types of both mobile and immobile Ca^2+^ buffers, for example, ATP, SERCA protein, and mitochondria, into one cytoplasmic Ca^2+^ buffering site (Shannon et al., 2004; Means et al., 2006; Nelson et al., 2014). *CER* and *KER* values were 3.6 and 2 mM, respectively (Cheung, 1980; Robertson et al., 1981; Means et al., 2006). For each transfectant, the *Amax*, *KACa*, and *KICa* were experimental values when *nA* is regarded as 2 (Table 3). The *∊*, *kleak*, *Vmax,ER*, *Km,ER*, *vin*, *Vmax,out*, and *Km,out* were set at 500, 0.025/s, 0.046 mM/s, 0.0003 mM, 0.0021 mM/s, 0.0067 mM/s, and 0.0004 mM, respectively. The set of these parameters was chosen so that the ER Ca^2+^ concentrations in mock, WT RyR2, and R2474S-mutant transfectants were in the appropriate range. The maximum concentrations and periodic times for *CER* in mock and all transfectants were obtained from their steady state reached by the model simulation. All simulations were performed with the package “deSolve” in R 3.6.3 (http://www.r-project.org/). The ODE solver selected was “lsoda,” which switches automatically between stiff and non-stiff methods depending on the problem.

**Table 3.** Parameters for cytosolic and ER Ca^2+^ binding sites.

| Fixed parameters | value | reference |
| --- | --- | --- |
| $V_{ratio}$ | 0.03 | (Shannon et al., 2004) |
| $C_{cyt}$(mM) | 11 | (Means et al., 2006; Nelson et al., 2014) |
| $K_{cyt}$(mM) | 0.3 | |
| $C_{ER}$(mM) | 3.6 | (Means et al., 2006) |
| $K_{ER}$(mM) | 2 | |

### Data analysis

Data are presented as the mean ± SD. Statistical comparisons were performed using Prism 8 (GraphPad Software, Inc., La Jolla, CA). Student’s *t*-test was used to compare two groups. *P* values < 0.05 were considered significant.

## Results

Stable tetracycline-inducible HEK293 cell lines expressing WT and mutant RyR2s (10 CPVT and 6 artificial mutants) were generated. The CPVT mutations are located at the N-terminal helical domain (V2321M, R2474S), central domain (D3638A, Q4201R), divergent region 1 (K4392R), and transmembrane domain (R4497C, K4751Q, H4762P, K4805R, and I4867M) (Fig. 1A and Table 1). Furthermore, artificial mutations were made by single alanine substitutions at the S4–S5 linker region, in which alanine substitutions differently affect the CICR activity depending on the position of the alpha helix in RyR1 (Murayama et al., 2011). Figure 1B shows the western blot data of WT and mutant RyR2s at 26 h after induction. WT and mutant RyR2 proteins were similarly expressed in HEK293 cells, with the exception of K4751Q, K4805R and L4757A (Fig. 1B), the expression levels of which were lower than those of the other mutants. The low expression levels may correlate with the high Ca^2+^ release activity, as described later.

**Figure 1.**
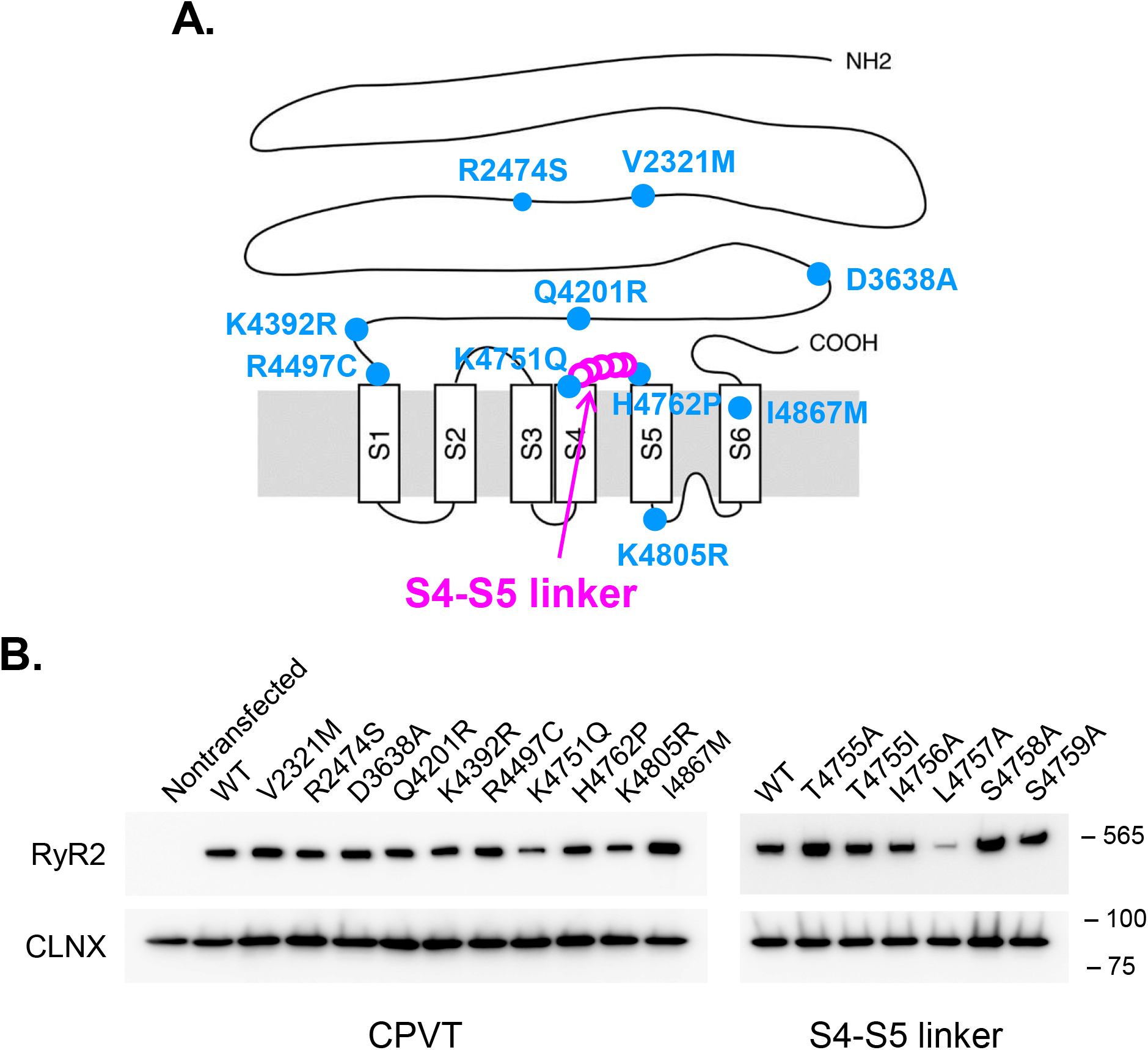
RyR2 mutations used in this study. **A.** Location of mutations in the primary structure of RyR2. CPVT mutations (blue circles) were based on human disorders (see Table 1), whereas the mutations in S4–S5 linker regions (pink open circles) were artificial. **B.** Western blot of WT and RyR2 mutants expressed in HEK293 cells. Note that the expression levels were similar to each other, with the exceptions of those of K4751Q, K4805R, and L4757A, which were significantly lower than the others. Calnexin (CLNX) was used as a loading control.

### Ca^2+^ homeostasis in HEK cells expressing WT and CPVT mutant RyR2s

Single-cell Ca^2+^ imaging in HEK cells was carried out 26–28 h after induction by doxycycline. Figure 2A shows representative cytoplasmic Ca^2+^ measurements obtained with fluo-4. HEK293 cells expressing WT-RyR2 showed spontaneous Ca^2+^ oscillations, as reported previously (Fig. 2A left) (Jiang et al., 2004; Fujii et al., 2017; Uehara et al., 2017). The application of 10 mM caffeine induced a transient Ca^2+^ release with a similar amplitude to spontaneous oscillations. Cells expressing R4497C- and R2474S-RyR2 showed Ca^2+^ oscillations with increased frequency and smaller amplitude compared with those of WT cells (Fig. 2A middle two). Most H4762P cells showed no clear oscillations and the application of caffeine caused only a small Ca^2+^ transient (Fig. 2A right). With the exceptions of H4762P and K4805R, more than 70% of WT and CPVT mutant cells showed clear Ca^2+^ oscillations. In H4762P and K4805R cells, only 10% of cells exhibited distinguishable Ca^2+^ oscillations whereas the remaining cells did not show clear oscillations. The average oscillation frequency was significantly higher in CPVT mutants than in WT, with the exception of K4392R (Fig. 2B). Average peak and resting cytosolic Ca^2+^ signals and calculated Ca^2+^ concentrations ([Ca^2+^]_cyt_) are shown in Supplemental Fig. 1 and Fig. 2C, respectively. Peak [Ca^2+^]_cyt_ was significantly smaller in CPVT than in WT, with the exception of K4392R. Resting [Ca^2+^]_cyt_ was significantly higher in K4751Q, H4762P, and K4805R cells (Fig. 2C, Supplemental Fig. 1B). A clear negative correlation was observed between the oscillation frequency and peak amplitude; that is, a higher frequency was associated with a smaller Ca^2+^ transient (Fig. 2D).

**Figure 2.**
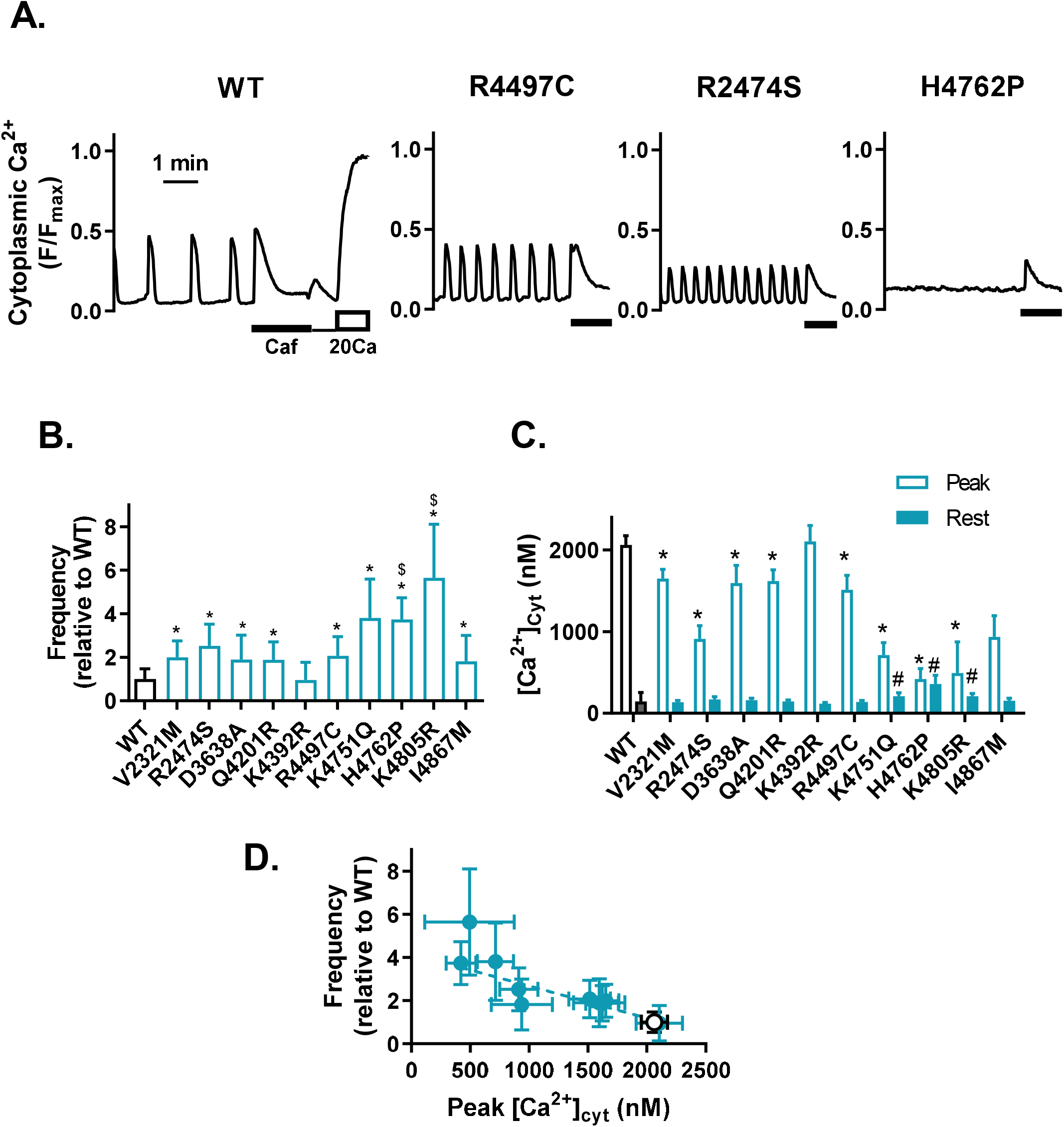
Cytoplasmic Ca^2+^ signals in HEK293 cells expressing WT and CPVT mutant RyR2s. **A.** Typical traces of fluo-4 Ca^2+^ signals in WT (left), R4497C (middle left), R2474S (middle right), and H4762P (right) cells. Fluorescence signals from fluo-4 (F) in individual cells were obtained in normal Krebs solution for 5 min, and then in the presence of 10 mM caffeine for 2 min (thick line). As shown in WT trace, at the end of each measurement, cells were treated with 0Ca-Krebs solution (thin line) and finally with 20Ca-Krebs solution (open bar), giving the maximal fluorescence intensity of fluo-4 (F_max_). Fluorescence signals are expressed as F/F_max_. **B.** Average oscillation frequencies in WT and CPVT cells. Oscillation frequency was determined by counting the number of Ca^2+^ oscillations that occurred during 5 min measurements. In contrast, for mutations marked with $, the average of only cells exhibiting clear oscillations is shown since only a small proportion of cells showed clear oscillations. Data are mean±SD. n=59–132, except data marked by $ (n=11–15), in which the proportion of oscillating cells was less than 10%. ^*^*p* < 0.05 compared with WT. **C.** Average peak and resting [Ca^2+^]_cyt_. Data are mean±SD. ^*^*p* < 0.05 compared with WT peak. ^#^*p* < 0.05 compared with WT rest, n=57–140. **D.** Relationship between oscillation frequency and peak [Ca^2+^]_cyt_. WT (black open circle), CPVT (blue filled circle). Data of H4762P and K4805R are not included.

We next measured ER Ca^2+^ ([Ca^2+^]_ER_) levels using R-CEPIA1er, a genetically encoded ER Ca^2+^ sensor protein (Suzuki et al., 2014; Murayama et al., 2015; Murayama et al., 2016; Fujii et al., 2017; Uehara et al., 2017). Figure 3A shows representative [Ca^2+^]_cyt_ and [Ca^2+^]_ER_ signals from G-GECO1.1 and R-CEPIA1er, respectively. HEK293 cells expressing WT RyR2 (Fig. 3A, left) showed a periodic decrease in R-CEPIA1er signals in normal Krebs solution, which was a reflection of Ca^2+^ release from the ER. Prior to each Ca^2+^ release, the R-CEPIA1er signal reached a maximal level (threshold), then rapidly decreased to reach a minimal level (nadir), and subsequently gradually increased again toward the threshold level. Both the threshold and the nadir levels were significantly reduced in R4497C, R2474S, and H4762P to varying degrees (Fig. 3A). Supplemental Fig. 2A and Fig. 3B indicate average ER Ca^2+^ signals and calculated [Ca^2+^]_ER_, respectively. All CPVT cells showed significantly decreased threshold and nadir [Ca^2+^]_ER_, with the exception of K4392R. There was a strong negative correlation between oscillation frequency and [Ca^2+^]_ER_ (Fig. 3C). A good correlation was also observed between nadir and threshold [Ca^2+^]_ER_ (Supplemental Fig. 2C).

**Figure 3.**
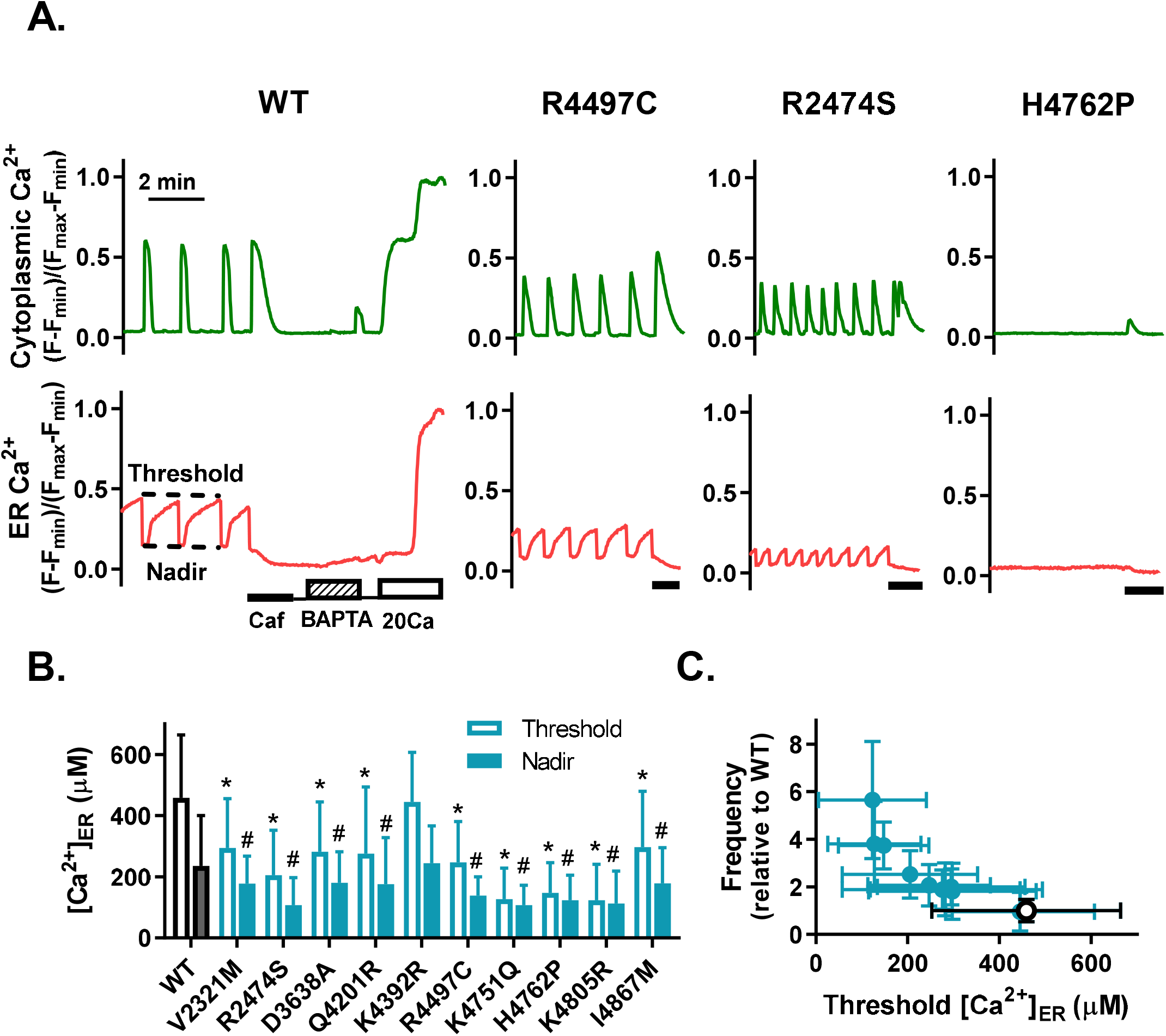
ER Ca^2+^ signals in HEK293 cells expressing WT and CPVT mutant RyR2s. **A.** Representative traces of G-GECO1.1 and R-CEPIA1er signals for WT (left), R4497C (middle left), R2474S (middle right), and H4762P (right). Ca^2+^ signals in individual cells were obtained in normal Krebs solution and then in caffeine solution (thick line). As shown in the WT traces, at the end of each measurement, the cells were perfused with 0Ca-Krebs solution (thin line), BAPTA-0Ca-Krebs solution (hatched bar), 0Ca-Krebs solution (thin line), and then 20Ca-Krebs solution (open bar). *Fmin* and *F*_max_ values were obtained with the BAPTA-0Ca-Krebs solution and 20Ca-Krebs solution, respectively. **B.** Average of threshold and nadir [Ca^2+^]_ER_ of cells expressing WT and CPVT mutants. Threshold [Ca^2+^]_ER_ was obtained from both oscillating and non-oscillating cells, while nadir [Ca^2+^]_ER_ was obtained only from oscillating cells. Data are mean±SD, ^*^*p* < 0.05 compared with WT threshold. ^#^*p* < 0.05 compared with WT nadir, n=57–130. **C.** Relationship between oscillation frequency and [Ca^2+^]_ER_ in cells expressing WT and CPVT mutant RyR2s. WT (open circle), CPVT (blue filled circle).

### Ca^2+^-dependent [^3^H]ryanodine binding activity

The properties of cytosolic Ca^2+^-dependent channel activity were then evaluated by [^3^H]ryanodine binding assay (Fig. 4). Both WT and mutant RyR2 channels exhibited bell-shaped biphasic Ca^2+^ dependence (Figure 4A) and the biphasic properties of CICR activity are effectively described by Equations (1)–(3) with three parameters: the gain (*A*_max_) and dissociation constants for activating Ca^2+^ (*KACa*) and inactivating Ca^2+^ (*KICa*) with fixed Hill coefficients (*n*_A_ = 2.0 and *n*_I_ = 1.0) (Murayama et al., 2000; Murayama and Kurebayashi, 2011) (Figure 4B). The [^3^H]ryanodine binding data for the mutant channels were subjected to the fitting of Equations (1)–(3) to obtain the three parameters (Table 2, Figure 4C–E). Six CPVT mutants, V2321M, R2474S, Q4201R, R4497C, K4751Q, and K4805R, had significantly increased *A*_max_, whereas four variants, D3638A, K4392R, H4762P, and I4867M, showed similar or slightly smaller *Amax* compared with WT. All CPVT mutants, with the exception of K4392R, showed significant increases of 1/*KA*. Among them, R2474S, K4751Q, H4762P, and K4805R showed marked increases of 1/*KACa* by 2-fold or more. *KICa* was increased in most mutants, with the exceptions of D3638A, K4392R, and I4867M. To compare Ca^2+^ release activity for WT and mutant RyR2s at resting [Ca^2+^]_cyt_, CICR activity at pCa7.0 (A_7.0_) was calculated using the three parameters (Table 2) and Equations (1)–(3), and expressed as a value normalized to that of WT (Fig. 4F). This model-based extrapolation procedure is very effective for evaluating CICR activities at low Ca^2+^, at which the direct determination of [^3^H]ryanodine binding is difficult (Murayama et al., 2015; Murayama et al., 2016; Nozaki et al., 2020). All mutations except K4392R enhanced the CICR activity by 2-fold or more. H4762P resulted in the highest CICR activity, followed by K4805R, K4751Q, and R2474S in this order. The threshold [Ca^2+^]_ER_ for Ca^2+^ release was plotted against A_7.0_ in log scale (Fig. 4G). There was a good inverse correlation between them, suggesting that the threshold [Ca^2+^]_ER_ is dependent on the CICR activity of RyR2. A strong correlation was also found between oscillation frequency and A_7.0_ (Fig. 4H).

**Figure 4.**
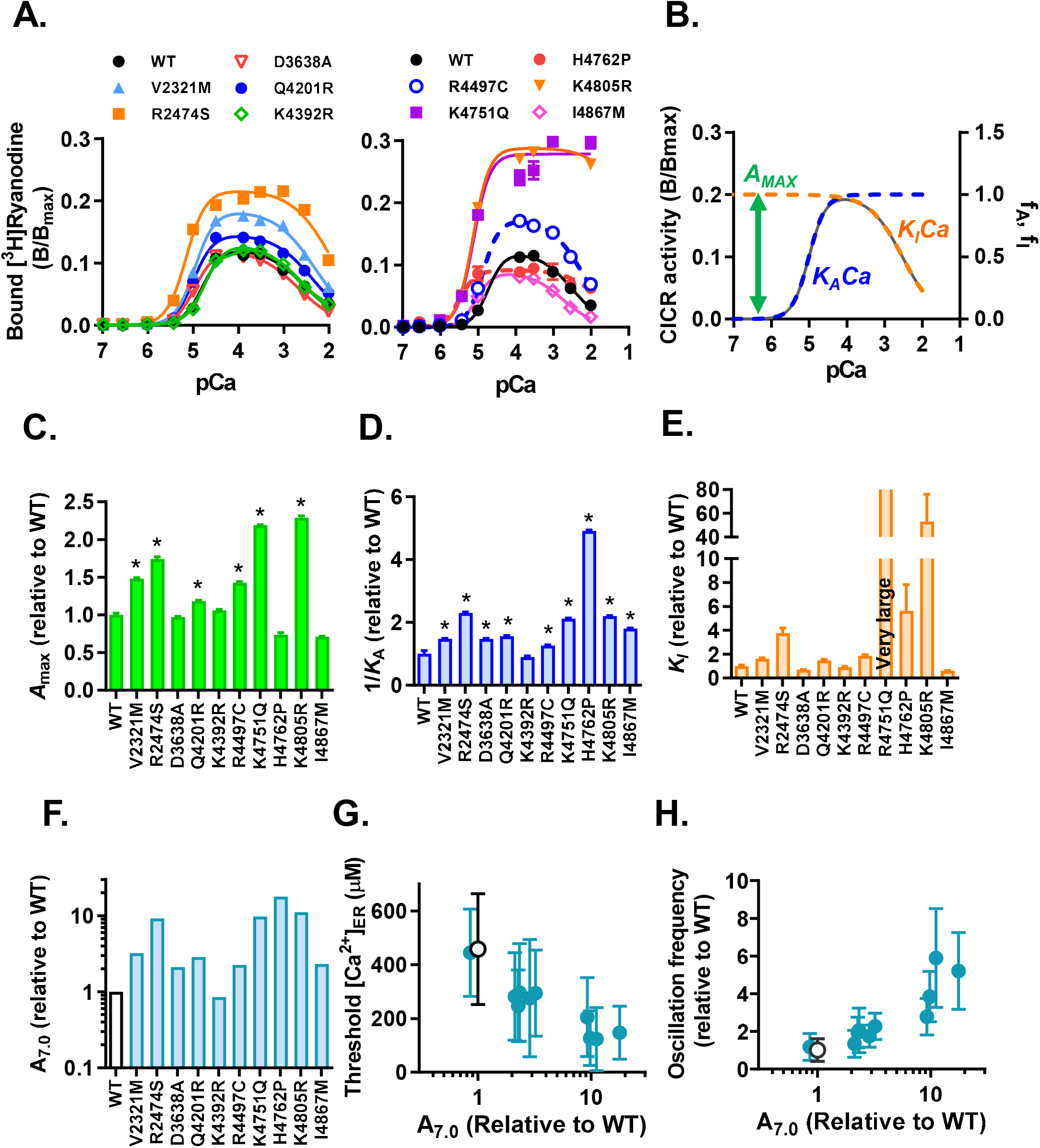
Ca^2+^-dependent [^3^H]ryanodine binding activity. **A.** Ca^2+^-dependent [^3^H]ryanodine binding activity of WT and CPVT mutants. **B.** The model of bell-shaped Ca^2+^ dependence composed of Eq. (1)–(3) with the three parameters: maximal activity (*Amax*), Ca^2+^ sensitivity for activation (*KACa*), and Ca^2+^ sensitivity for inactivating Ca^2+^ (*KIca*) (see Materials and Methods). **C–E.** Comparison of the three parameters, *A*_max_ (**C**), 1/*K*_ACa_ (**D**), and *K*_Ica_ (**E**), among WT and CPVT mutants. The three parameters were plotted as values relative to WT. Data are mean±SD, n = 3. ^*^*p* < 0.05 compared with WT. **F.** Calculated CICR activities at resting [Ca^2+^]_cyt_ (A_7.0_) of CPVT mutants normalized to that of WT. **G.** Relationship between threshold [Ca^2+^]_ER_ and A_7.0_. **H.** Relationship between oscillation frequency and A_7.0_.

### Effects of artificial mutations on CICR activity and [Ca^2+^]_ER_

The above results indicate that Ca^2+^ oscillation frequency and [Ca^2+^]_ER_ correlate well with the CICR activity at resting [Ca^2+^]_cyt_. We next examined whether this is also the case for artificial mutations associated with various CICR activities. We previously reported that single alanine substitutions at the S4–S5 linker region in RyR1 differently affect the three parameters of CICR activity depending on the position of the alpha helix (Murayama et al., 2011). On the basis of a high degree of sequence homology between RyR1 and RyR2 at the S4–S5 region, the homologous mutations were made in RyR2 and their effects on [Ca^2+^]_ER_ were examined (Fig. 5A). The [^3^H]ryanodine binding assay revealed that the S4–S5 linker region mutations had different degrees of influence on the CICR activity, A_7.0_, depending on the mutation site (Fig. 5A). A_7.0_ was greatly enhanced by L4756A (82-fold) and T4754I (9-fold), moderately enhanced by I4755A and S4757A (2-fold), but significantly suppressed by T4754A (0.6-fold) and S2758A (0.3-fold) (Fig. 5B). These changes by individual mutations were similar to those at corresponding sites in RyR1 (Murayama et al., 2011).

**Figure 5.**
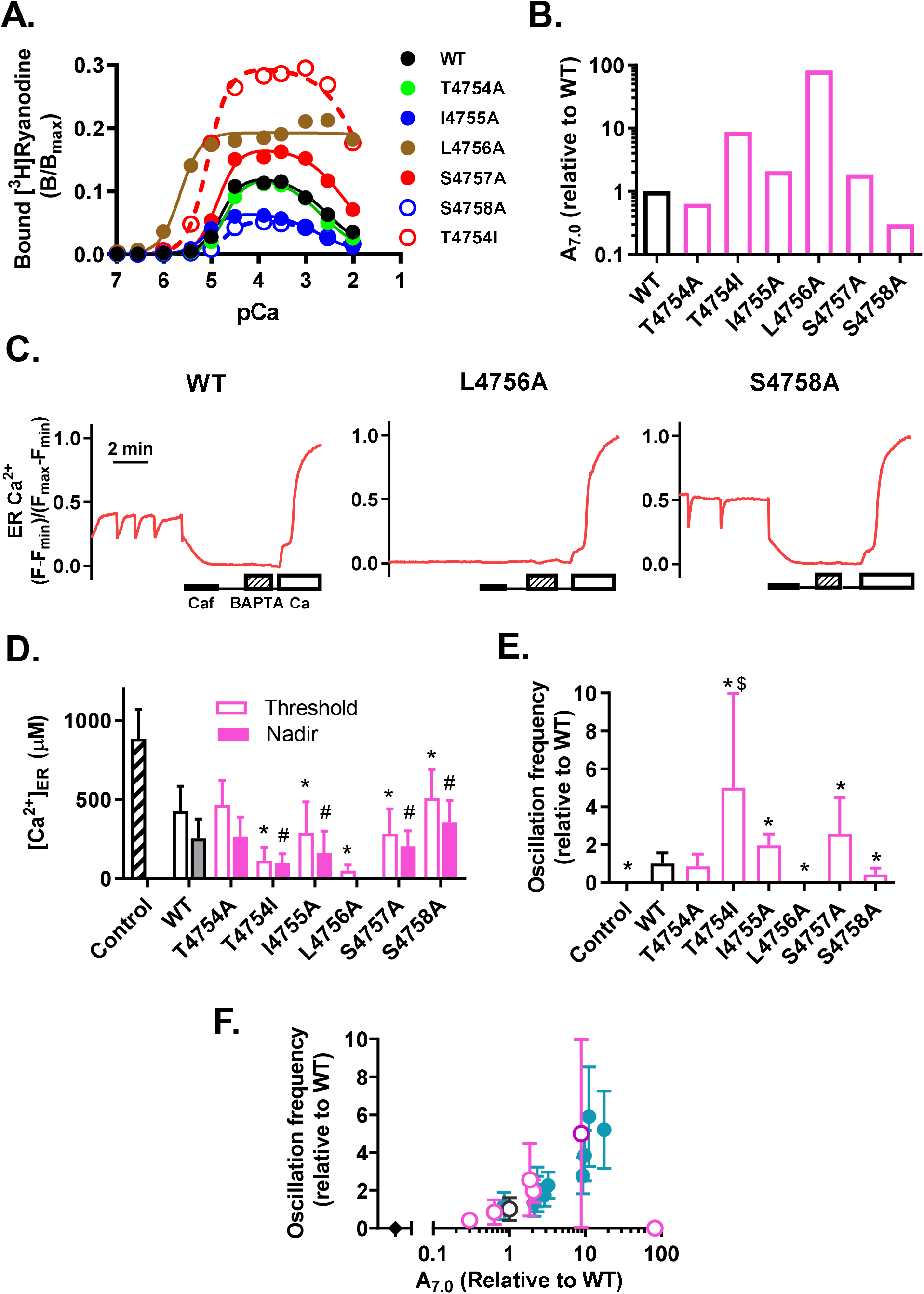
Effects of artificial mutations at S4–S5 linker region on Ca^2+^-dependent [^3^H]ryanodine binding activity and Ca^2+^ homeostasis in HEK293 cells. **A.** Ca^2+^-dependent [^3^H]ryanodine binding activity of WT and artificial mutants. **B.** Calculated relative A_7.0_ based on the [^3^H]ryanodine binding data. **C.** Representative [Ca^2+^]_ER_ of WT, L4756A, and S4758A. ER Ca^2+^ signals in individual cells were obtained in normal Krebs solution and then in caffeine solution (thick line). At the end of each measurement, the cells were perfused with 0Ca-Krebs solution (thin line), BAPTA-0Ca-Krebs solution (hatched bar), 0Ca-Krebs solution (thin line), and then 20Ca-Krebs solution (open bar). **D.** Average threshold and nadir [Ca^2+^]_ER_ levels. Threshold [Ca^2+^]_ER_ was obtained from both oscillating and non-oscillating cells, while nadir [Ca^2+^]_ER_ was obtained only from oscillating cells. Only threshold [Ca^2+^]_ER_ is shown for control HEK293 and L4756A cells because they did not show Ca^2+^ oscillation. Data are mean±SD. n=30–86. ^*^*p* < 0.05 compared with WT threshold. ^#^*p* < 0.05 compared with WT nadir. **E.** Average oscillation frequency of the S4–S5 linker region mutants. Data are mean±SD (n=41–60) except for T4754I (n=19), in which the proportion of oscillating cells was 30% and average frequency was obtained from only cells showing Ca^2+^ oscillations. ^*^*p* < 0.05 compared with WT. **F.** Relationship between oscillation frequency and A_7.0_. S4–S5 linker mutants (pink open circle), control HEK (filled diamond), WT (open circle), and CPVT (blue filled circle).

We then determined [Ca^2+^]_ER_ in HEK293 cells. L4756A showed almost complete depletion of [Ca^2+^]_ER_ and no Ca^2+^ oscillations (Fig. 5C middle, 5D, 5E). T4754I, I4755A, and S4757A also significantly reduced the threshold and nadir of [Ca^2+^]_ER_. In contrast, S4758A showed higher [Ca^2+^]_ER_ than WT, but still lower than that in control HEK293 cells (Fig. 5C right and Fig. 5D, Fig. 2B). No significant difference was found between T4754A and WT. T4754I, I 4755A, and S4757A showed more frequent Ca^2+^ oscillations whereas S4758A exhibited less frequent oscillations than WT. There was also a good correlation between CICR activity and Ca^2+^ oscillation frequency (Fig. 5F); that is, mutants with higher CICR activity showed more frequent Ca^2+^ oscillation, although a mutant with extremely high activity, L4756A, did not show any oscillations because of ER Ca^2+^ depletion.

Taking the data of CPVT and S4–S5 mutants together, we analyzed the relationships between threshold [Ca^2+^]_ER_ and CICR parameters (Fig. 6). The threshold [Ca^2+^]_ER_ was significantly correlated with *Amax* (R^2^=0.136, P<0.0001), 1/*KACa* (R^2^=0.726, P<0.0001), and *KICa* (R^2^=0.504, P=0.0021) (Fig. 6A–C). However, the goodness-of-fit statistic was the best for [Ca^2+^]_ER_ vs. A_7.0_ (R^2^=0.869, P<0.0001) (Fig. 6D). Taken together, these findings indicate that threshold [Ca^2+^]_ER_ and oscillation frequency are highly dependent on A_7.0_, the CICR activity of RyR2 at resting [Ca^2+^]_cyt_.

**Figure 6.**
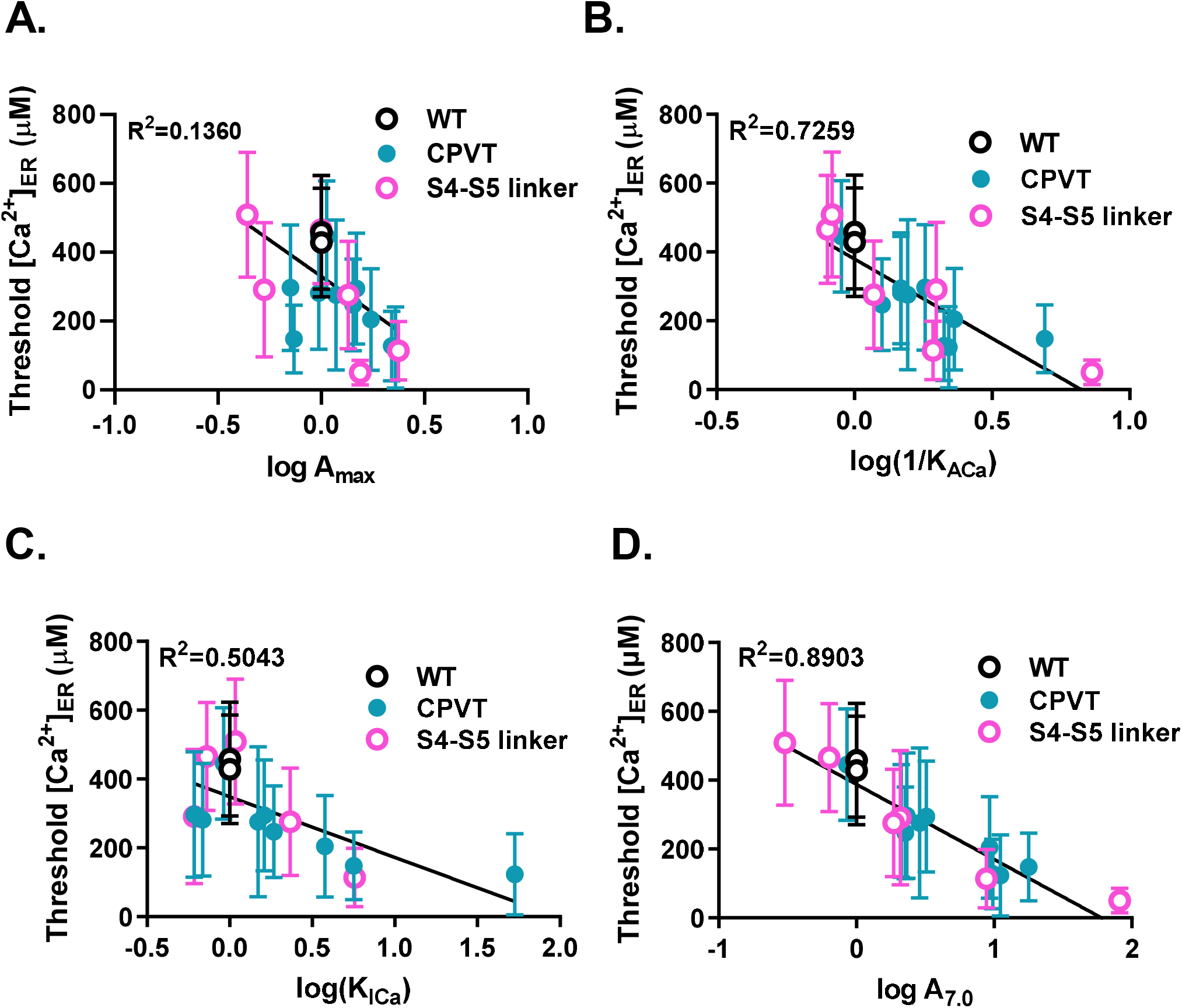
Comparison of goodness of fit between threshold [Ca^2+^]ER and CICR parameters. **A.** [Ca^2+^]_ER_ vs log*A7.0*. **B.** [Ca^2+^]_ER_ vs *Amax*. **C.** [Ca^2+^]_ER_ vs. log(1/*KACa*). **D.** [Ca^2+^]_ER_ vs. log(*KICa*).

### Effects of expression level of RyR2 on Ca^2+^ homeostasis

We previously showed that the threshold [Ca^2+^]_ER_ for Ca^2+^ release decreases with time after the induction of RyR2 expression in HEK293 cells (Uehara et al., 2017). This implies that the threshold [Ca^2+^]_ER_ is not fixed but affected by the expression levels of RyR2. We examined the relationship between threshold [Ca^2+^]_ER_ and the RyR2 expression level using WT and R2474S. The protein expression of WT and R2474S increased with time after induction (Fig. 7A) and reached a quasi-steady state at around 24 h (Fig. 7B). The threshold [Ca^2+^]_ER_ in both WT and R2474S cells decreased with time, and that in R2474S cells always decreased faster than that in WT cells (Fig. 7C, D). As a consequence, threshold [Ca^2+^]_ER_ in R2474S cells was lower than that in WT at the same expression level (Fig. 7E). On the basis of these findings, we hypothesized that the net Ca^2+^ release rates *in situ* in RyR2-expressing cells are proportional to the product of the number of RyR2 channels and their own CICR activity at resting [Ca^2+^]_cyt_, A_7.0_. There was a clear correlation between the threshold [Ca^2+^]_ER_ and the net Ca^2+^ release rate, regardless of the type of mutant (Fig. 7F). These results support the idea that threshold [Ca^2+^]_ER_ is determined by both the CICR activity at resting [Ca^2+^]_cyt_ and the density of RyR2 channels on the ER membrane.

**Figure 7.**
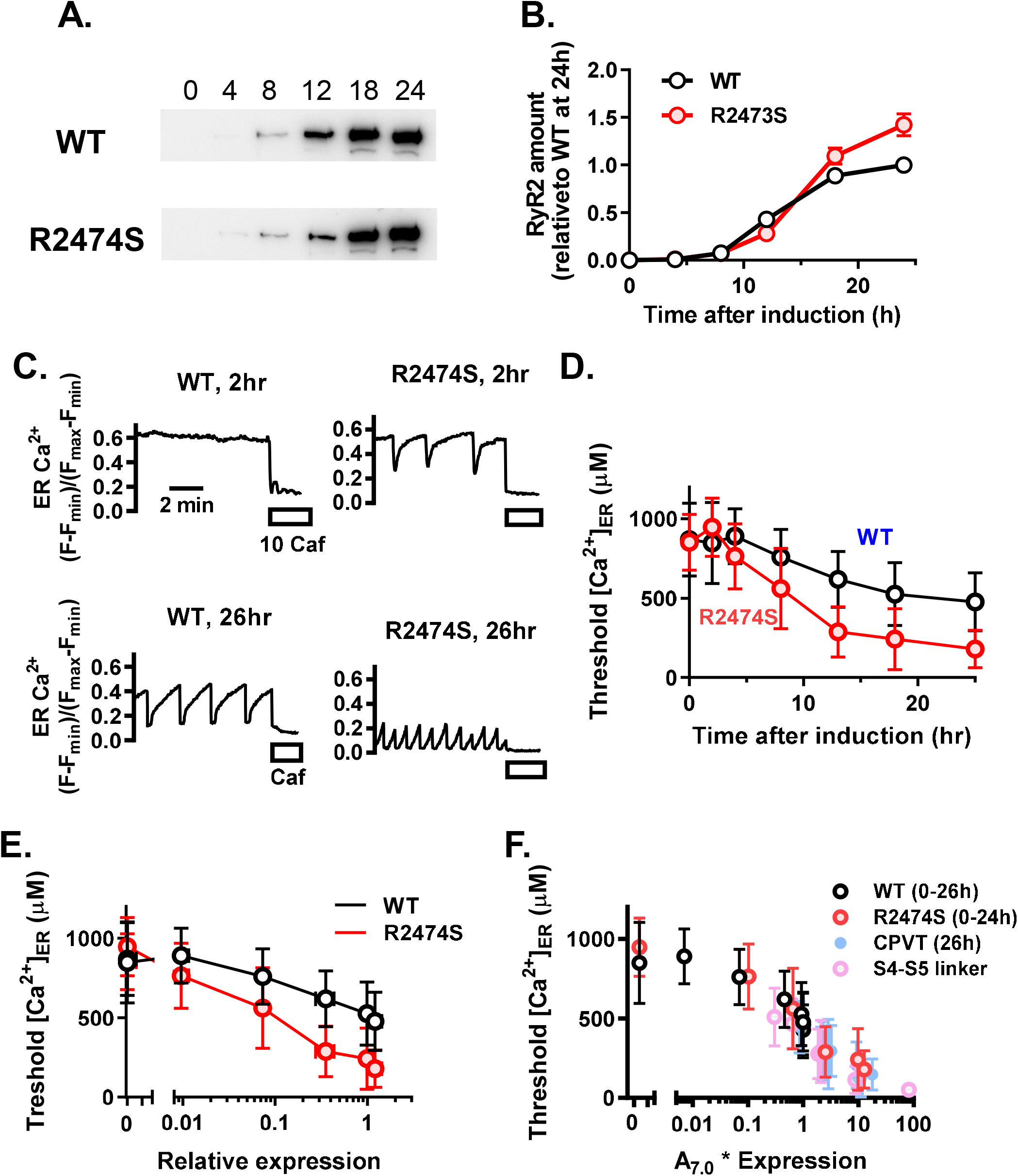
Correlation between threshold [Ca^2+^]ER and expression level of RyR2. **A.** Western blotting analysis of expression levels for WT and R2474S at 0, 4, 8, 12, 18, and 24 h after induction. **B.** Time course of expression level after induction. **C.** Typical ER Ca^2+^ signals at 2 and 26 h after induction. **D.** Time course of threshold [Ca^2+^]_ER_ after induction. **E.** Relationship between threshold [Ca^2+^]_ER_ and expression level. **F.** Correlation between threshold [Ca^2+^]_ER_ and net CICR activity at resting [Ca^2+^]_cyt_, a product of A_7.0_ and expression.

### Simulation of cellular Ca^2+^ dynamics using mathematical model

The strong correlation between threshold [Ca^2+^]_ER_ and CICR activity at resting [Ca^2+^]_cyt_, A_7.0_, suggests that threshold [Ca^2+^]_ER_ may be mainly determined by CICR activity. We tested the plausibility of this hypothesis by using a mathematical model. To mathematically describe calcium dynamics, cells were assumed to comprise cytosolic and ER compartments (Fig. 8A), where the transport of Ca^2+^ between the cytosol and the ER involves [Ca^2+^]_cyt_-dependent Ca^2+^ release through RyR2, saturable ER Ca^2+^ uptake by SERCA, and a passive leakage pathway that is intrinsic to HEK293 cells. Cellular Ca^2+^ influx via the plasma membrane is assumed to obey zero-order kinetics provided that the extracellular bulk concentration is constant and Ca^2+^ efflux is driven by a presumptive Ca^2+^ pump. In actual cells, it is necessary to consider the effect of diffusion, but here, for the sake of simplicity, we assumed a rapid equilibrium of mobilized Ca^2+^. At first, we tested whether WT Ca^2+^ oscillation is reproduced by the model. For each mutant, the *Amax*, *KACa*, and *KICa* were experimental values when *nA* and *nI* are regarded as 2 and 1, respectively (Table 2). The *Vratio*, *Ccyt*, *Kcyt*, *CER*, and *KER* values are shown in Table 3. The *∊*, *kleak*, *Vmax,ER*, *Km,ER*, *vin*, *Vmax,out*, and *Km,out* were set at 500, 0.025/s, 0.046 mM/s, 0.0003 mM, 0.0021 mM/s, 0.0067 mM/s, and 0.0004 mM, respectively (Figure 8A right), to satisfy the following three conditions: (1) in mock HEK293 cells, Ca^2+^ concentration in ER at a steady state was 0.6−0.8 mM; (2) in WT RyR2 cells, Ca^2+^ maximum concentration in ER was 0.3−0.5 mM and periodic time was about 60 s; and (3) in R2474S mutant, Ca^2+^ maximum concentration in ER was 0.1−0.2 mM. The mathematical model could reproduce the periodic cytoplasmic Ca^2+^ increase and [Ca^2+^]_ER_ decrease, which were similar to the [Ca^2+^]_cyt_ and [Ca^2+^]_ER_ observed in WT, and no oscillation in control HEK (Fig. 8B, second from the left). Then, using the same constants and Ca^2+^ release activity with the three parameters for individual mutants, Ca^2+^ oscillations for mutant cells were calculated. The model also effectively reproduced a reduced threshold [Ca^2+^]_ER_ for Ca^2+^ release and an increased oscillation frequency in R4497C and R2474S (Fig. 8B, middle), and almost completely diminished Ca^2+^ oscillation with a further decrease in [Ca^2+^]_ER_ in H4762P cells (Fig. 8B, right). Good correlations of simulated threshold [Ca^2+^]_ER_ vs. A_7.0_ and simulated oscillation frequency vs. A_7.0_ were reproduced in the A_7.0_ range of 0.3–10, as shown in Fig. 8C and D. Furthermore, the lack of oscillation in cells with very high CICR activity, L4756A, was also reproduced (Fig. 8C and D, rightmost point). Figure 8E shows the relationship between actual and calculated [Ca^2+^]_ER_. The simulated [Ca^2+^]_ER_ for mutant RyR2s effectively recapitulated the actual cytoplasmic and [Ca^2+^]_ER_ signals. These results strongly support the idea that [Ca^2+^]_ER_ is largely determined by the CICR activity at resting [Ca^2+^]_cyt_.

**Figure 8.**
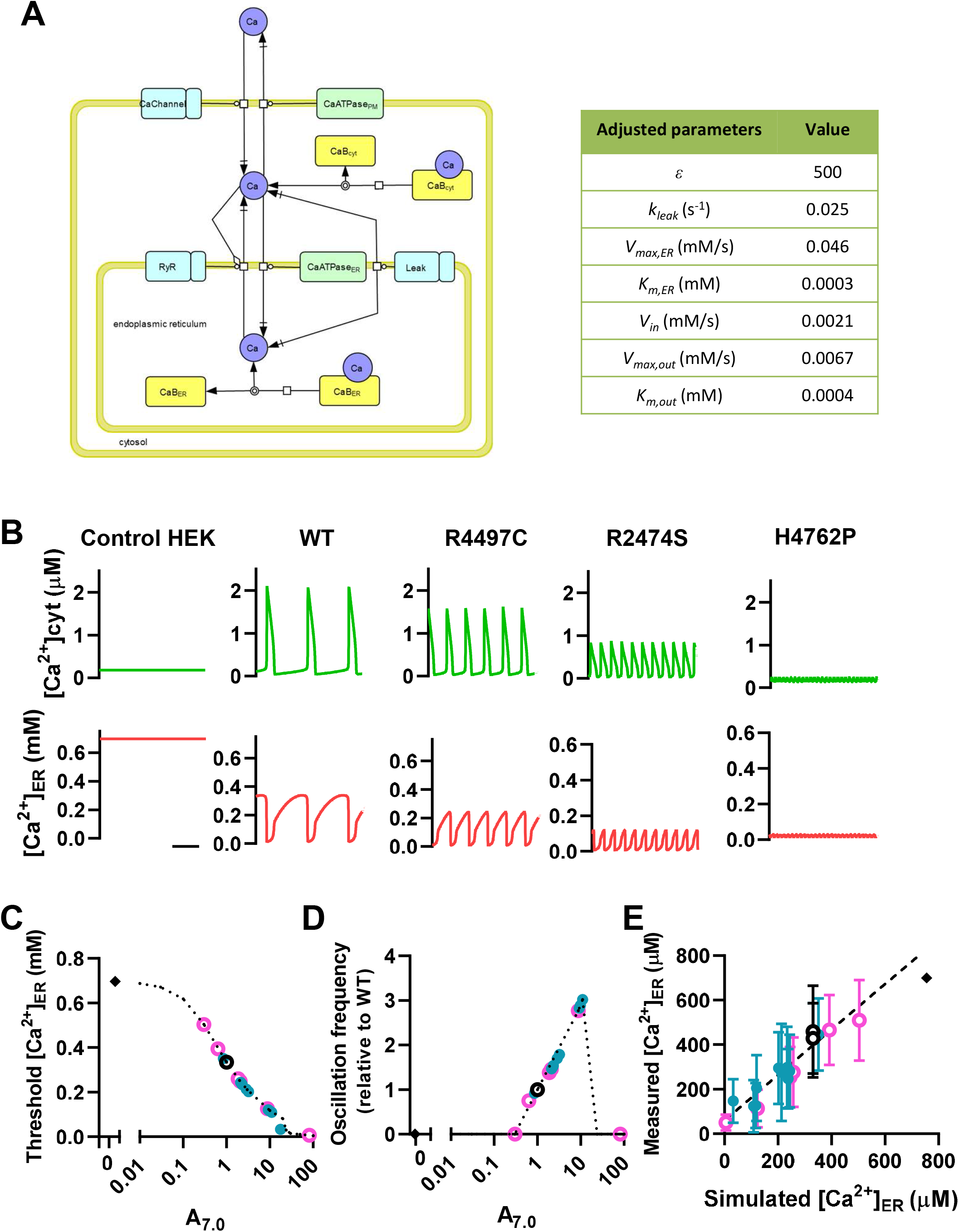
Mathematical model of Ca^2+^ homeostasis based on CICR activity. **A.** Ca^2+^ dynamics model (left) and adjusted parameters for the simulation (right). **B.** Typical traces of simulated [Ca^2+^]_cyt_ and [Ca^2+^]_ER_ in Control HEK, WT, R4497C, R2474S, and H4762P. **C.** Relationship between simulated threshold [Ca^2+^]_ER_ and CICR activity at resting [Ca^2+^]_Cyt_, A_7.0_, for WT and mutant RyR2s. Dotted line indicates the simulated relationship in the range of A_7.0_ from 0 to 100. Note that the relationship is close to a straight line in the A_0_ range of 0.3–10. **D.** Relationship between simulated oscillation frequency and A_7.0_. **E.** Relationship between measured and simulated threshold [Ca^2+^]_ER_. Control HEK (filled diamond), WT (open circle), CPVT (blue filled circle), and S4–S5 linker mutants (pink open circle).

### Pathological relevance of CICR activity in CPVT mutations

The results in Fig. 2–4 indicate that the CICR activities at resting [Ca^2+^]_cyt_, A_7.0_, of CPVT mutants examined in this study ranged from mild to severe, with ~10-fold difference in the activities. To determine the pathological relevance of CICR activity in CPVT mutations, the age of onset of arrhythmia or SCD was plotted against the CICR activity (Fig. 9). Among the mutations with high CICR activity, R2474S, K4751Q, and K4805R caused disease at a young age, less than 8 years old, and only one or two (twin) individuals were affected in the family. In contrast, two of the mild mutations, Q4201R and R4497C, were detected in multiple individuals with various ages of onset from young to old age, indicating relatively high penetrance (Laitinen et al., 2001; Priori et al., 2001). Only a single case has been reported for each of the other mutations with moderate CICR activity, V2321M, D3638A, and I4867M. The H4762P proband carries another mutation, G4662S, and her three family members with the H4762P mutation have been reported to be asymptomatic. Notably, a patient with K4392R did not show any difference from WT. The mother of this patient bears the same mutation and is asymptomatic (Arakawa et al., 2015). After the publication of this report, K4392R (rs753733164) was found to be a benign variant that does not exert arrhythmogenic effects in a majority of its carriers.

**Figure 9.**
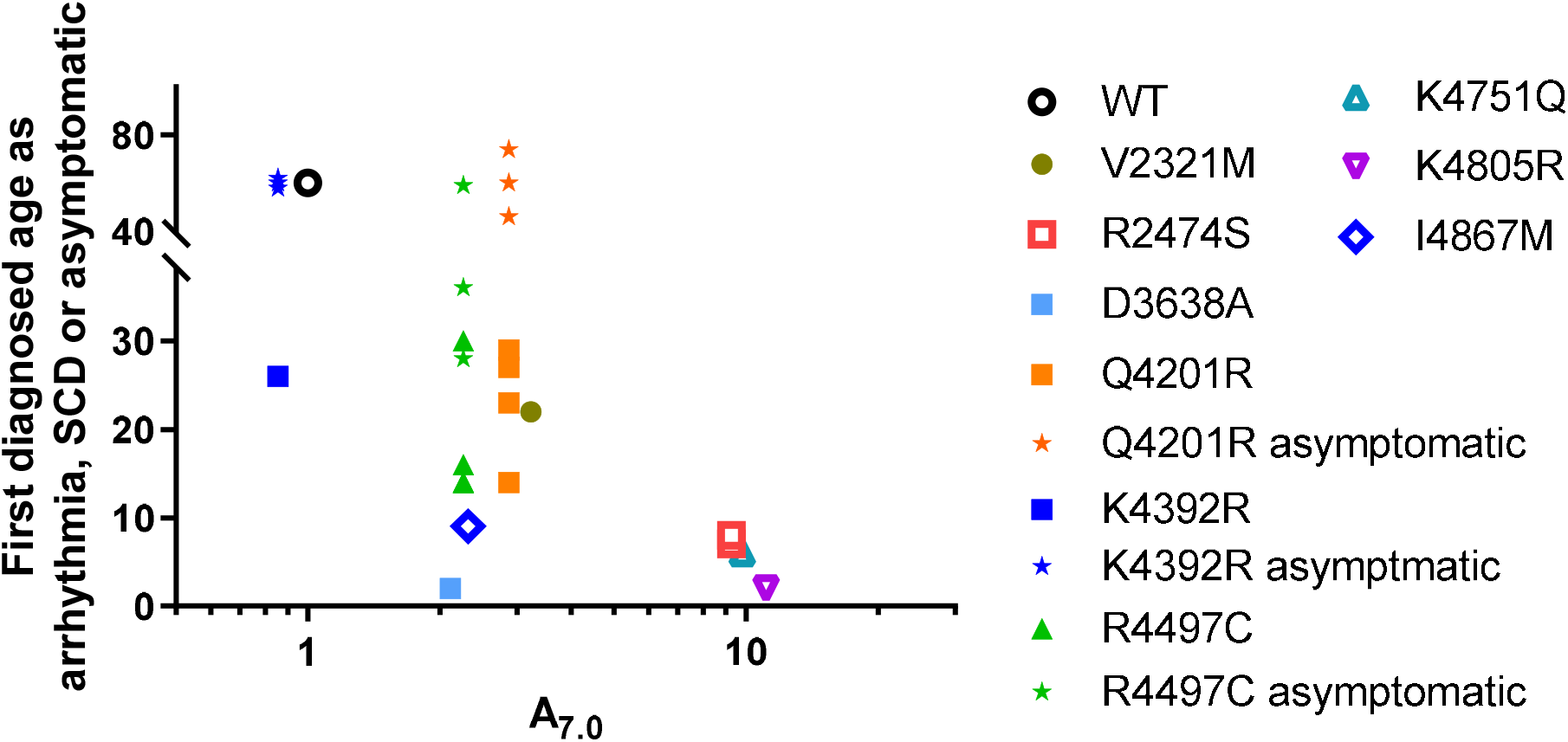
Relationship between the age of onset of arrhythmic symptom or sudden cardiac death and relative CICR activity of CPVT mutants. Individual points indicate individual RyR2 mutation carriers. Note that some carriers of Q4201R and R4497C were asymptomatic even in old age. The data for the H4762P patient are not plotted because she has two mutations.

## Discussion

The CPVT mutations in RyR2 are prone to cause spontaneous Ca^2+^ release, which leads to lethal arrhythmia. The HEK293 expression system is a good tool for studying the properties of mutant RyR2s and their effects on Ca^2+^ homeostasis. We measured [Ca^2+^]_cyt_ and [Ca^2+^]_ER_ in cells expressing WT and mutant RyR2s and determined CICR activity with [^3^H]ryanodine binding assay to explore the mechanisms controlling spontaneous Ca^2+^ release and the determinants of the threshold [Ca^2+^]_ER_ for Ca^2+^ release. Our quantitative analysis indicates that the CPVT mutations are associated with enhanced CICR activity and enable ranking of the severity of these mutations. Importantly, the CICR activity of RyR2 at resting [Ca^2+^]_cyt_ is critically involved in determining the threshold [Ca^2+^]_ER_ for spontaneous Ca^2+^ release.

### CICR activity at resting [Ca^2+^]_cyt_ is enhanced in all CPVT mutants

One of the major differences between this and previous studies is the Ca^2+^-dependent [^3^H]ryanodine binding activity of the mutant RyR2. For example, R2474S and R4497C mutations did not significantly affect [^3^H]ryanodine binding in the studies by Jiang et al. (Jiang et al., 2004; Jiang et al., 2005), whereas these mutations increased CICR activity at resting [Ca^2+^]_cyt_, A_7.0_, by 9.25- and 2.26-fold, respectively, in this study (Fig. 4).

There are two possible reasons for this difference. One is in the data analysis of [^3^H]ryanodine binding. We expressed the binding activity as B/B_max_, in which the determined activity was normalized by the number of total RyR2 molecules in the preparation (Murayama and Kurebayashi, 2011; Murayama et al., 2015; Fujii et al., 2017; Uehara et al., 2017) (see Methods). With this method, we show that most CPVT mutants including R2474S and R4497C significantly increased the *A*_max_ (Fig. 4). In contrast, in many studies, the binding activity was normalized by the peak values at the optimal Ca^2+^ (pCa ~4) (Jiang et al., 2004; Jiang et al., 2005; Xiao et al., 2016). This will mask the difference in *Amax*. Second, our fitting and extrapolation procedure using Eq. (1)–(3) makes quantitative differences in CICR activity at resting [Ca^2+^]_cyt_ clearer, at which the direct determination of [^3^H]ryanodine binding activity is difficult. In particular, since the parameter *KACa* contributes to CICR activity with a squared weight, the determination of *KACa* by a fitting procedure is important. In fact, in the study by Jiang et al. (2005), R2474S appeared to have higher *KACa* than WT. Although the activities at resting [Ca^2+^]_cyt_ are very low compared with the maximum value, they are not zero and certainly cause Ca^2+^ release to affect basal [Ca^2+^]_ER_. A similar conclusion was obtained with HEK293 cells expressing RyR1 (Murayama et al., 2015; Murayama et al., 2016; Murayama et al., 2018). Consequently, all CPVT mutations increase CICR activity at resting [Ca^2+^]_cyt_, although the mechanism of activation is mutation-site-specific; some substantially increase sensitivity for *KACa* whereas others predominantly increase *Amax*.

### CICR activity at resting [Ca^2+^]_cyt_ mainly determines threshold [Ca^2+^]_ER_

The development of ER-targeted Ca^2+^ indicator proteins with low Ca^2+^ affinity enabled us to determine [Ca^2+^]_ER_. R-CEPIA1er has an adequate K_D_ (565 µM) for the determination of [Ca^2+^]_ER_ in cells expressing RyRs (Suzuki et al., 2014; Bovo et al., 2016; Fujii et al., 2017; Uehara et al., 2017) compared with Fret-based ER-targeted indicator D1er (K_D_≈60 µM). Quantitative [Ca^2+^]_ER_ measurements were performed by the determination of F_max_ and F_min_ using high-Ca^2+^ and BAPTA–Krebs solutions containing a high concentration of Ca^2+^-ionophore at the end of the experiments.

Many previous reports indicated that HEK293 cells expressing RyR2 with CPVT mutation show reduced threshold [Ca^2+^]_ER_ for spontaneous Ca^2+^ release (Jiang et al., 2004; Jiang et al., 2005; Xiao et al., 2016). This study confirmed the results in a more quantitative manner. However, there are two possible explanations for this phenomenon: [Ca^2+^]_ER_ is determined either by CICR activity, the regulation by Ca^2+^ on the cytoplasmic side, or by the SOICR mechanism, the regulation by Ca^2+^ on the ER luminal side. Our results indicate that threshold [Ca^2+^]_ER_ for Ca^2+^ release for individual mutants varies depending on the net permeability of Ca^2+^ through RyR2 channels. Our study revealed that (1) with the same mutant, higher expression of RyR2 channels lowers the threshold [Ca^2+^]_ER_; (2) at the same expression level, RyR2 mutant with higher CICR activity presents a lower threshold [Ca^2+^]_ER_; and (3) the [Ca^2+^]_ER_ inversely correlates with net CICR activity at rest, as a product of mutant-specific CICR activity at rest and density of RyR2 molecules in the cell, probably on the ER membrane.

### Mathematical model for Ca^2+^ handling in cells expressing RyR2

We simulated Ca^2+^ influx and efflux across the cytoplasmic and ER membranes in HEK293 cells using a minimal essential model instead of detailed models with many components (Fig. 8). In the simulation, Ca^2+^ oscillation was successfully reproduced when the CICR parameter and the Ca^2+^ flux parameters were well balanced. Although Ca^2+^ at the luminal side activates RyR2 in lipid bilayer experiments and Ca^2+^ measurements in cardiac myocytes (Lukyanenko et al., 1996; Satoh et al., 1997; Sitsapesan and Williams, 1997), this simple model reasonably reproduced the Ca^2+^ oscillations by RyR2 and the effects of changes in CICR activity on Ca^2+^ oscillation frequency, threshold [Ca^2+^]_ER_ level, and peak amplitude of [Ca^2+^]_cyt_ without considering the effect of luminal Ca^2+^ activation. This may be due to the fact that HEK293 cells do not have cardiac-specific luminal proteins such as calsequestrin and the [Ca^2+^]_ER_ is less than 1 mM.

There were two non-negligible problems that this simulation did not reproduce. One is that resting [Ca^2+^]_cyt_ was not increased by CPVT mutants with high CICR activity. This is expected to be solved by adding store-operated Ca^2+^ (SOC) influx by [Ca^2+^]_ER_ depletion. Furthermore, our model did not reproduce mutant-dependent nadir [Ca^2+^]_ER_ levels. This may be improved by the reduction in Hill coefficient (*nA*) of the Ca^2+^-dependent activation of RyR2. Additionally, the consideration of SERCA activity depending on [Ca^2+^]_ER_, that is, lowered [Ca^2+^]_ER_ facilitating Ca^2+^ uptake, may also improve the simulation.

A general understanding of the overall influx and efflux balance is crucial in considering the likelihood of Ca^2+^ oscillations through RyR2, which are closely associated with the origin of arrhythmias. Importantly, our simulation indicates that the effects of CPVT mutations on the basic phenomenon of cytoplasmic Ca^2+^ oscillations and corresponding [Ca^2+^]_ER_ changes can be explained only by the difference in CICR activity at resting [Ca^2+^]_cyt_ determined by [^3^H]ryanodine binding measurements.

### CICR activity is a useful parameter in predicting the severity of CPVT mutations

Next-generation sequencing technology has revealed numerous novel RyR2 mutations linked to arrhythmias, but we cannot deduce their functional effects from the results of such sequencing. A heterologous expression system with HEK293 cells has been successfully used to assess the phenotypes resulting from mutations in RyR2 because they include loss-of-function and gain-of-function mutations (Jiang et al., 2002; Jones et al., 2008; Fujii et al., 2017; Uehara et al., 2017). We here show that the CICR activity of mutant RyR2s correlates well with the age of onset of arrhythmia or SCD, an index of severity of the disease (Fig. 9), although SCD can occur even with mutations with mild effects. Similar prediction of severity has been successfully performed for RyR1-carrying malignant hyperthermia (MH) and central core disease (CCD) mutations (Murayama et al., 2015; Murayama et al., 2016). Quantitative analysis of the CICR activity will be useful not only to distinguish the phenotypes but also to assess the severity of diseases.

### Limitations

This study investigated the properties of the homotetrameric mutant channels expressed in HEK293 cells. Because CPVT patients have heterozygous RyR2 channels, their changes in activity are expected to be smaller than those of homozygous channels. Therefore, the [Ca^2+^]_ER_ in cells having heterozygous channels may not be reduced as much as that in cells having homozygous ones. Nonetheless, we need to determine the activities of heterotetrameric channels consisting of WT and mutant RyR2 with various combinations. The second limitation is that we do not know the effects of mutations on interactions between RyR2 and cardiomyocyte-specific proteins such as FKBP12.6, traidin, calsequestrin, PKA, and CaMKII because we only performed observations in HEK293 cells. Furthermore, other genetic and environmental factors are also considered to be involved in human diseases. However, clarifying the impact of mutations themselves on RyR2 channel activity provides an essential basis for understanding the mutant RyR2 channel regulation and for designing therapeutic strategies.

## Acknowledgments

We thank Ikue Hiraga and the Laboratory of Radioisotope Research, Research Support Center, Juntendo University Graduate School of Medicine, for technical assistance. We also thank Edanz (https://en-author-services.edanz.com/ac) for editing the English text of a draft of this manuscript. This work was supported by JSPS KAKENHI Grant Number 19K07105 to N.K., 19H03404 to T.M., the Practical Research Project for Rare/Intractable Diseases (19ek0109202 to S.O. and N.K.) from the Japan Agency for Medical Research and Development (AMED), Platform Project for Supporting Drug Discovery and Life Science Research (Basis for Supporting Innovative Drug Discovery and Life Science Research (BINDS) (JP19am0101080 to T.M.), an Intramural Research Grant (2-5) for Neurological and Psychiatric Disorders from the National Center of Neurology and Psychiatry to T.M., and the Vehicle Racing Commemorative Foundation to T.M.

## Author contributions

N. Kurebayashi and T. Murayama carried out the cell biological experiments; R. Ohta and F Yamashita developed the simulation program; R. Ohta, F Yamashita and N Kurebayashi performed simulation and analysis; J. Suzuki, K. Kanemaru and M. Iino provided experimental tools; N. Kurebayashi, T. Murayama, R. Ohta, J. Suzuki, K Kanemaru and T. Sakurai analyzed the experimental data; S. Ohno and M. Horie analyzed the clinical data; and N. Kurebayashi, T Murayama, S. Ohno, M. Iino and F. Yamashita wrote the manuscript. All authors discussed the results and approved the final version of the manuscript.

---

**Supplemental Figure 1.**
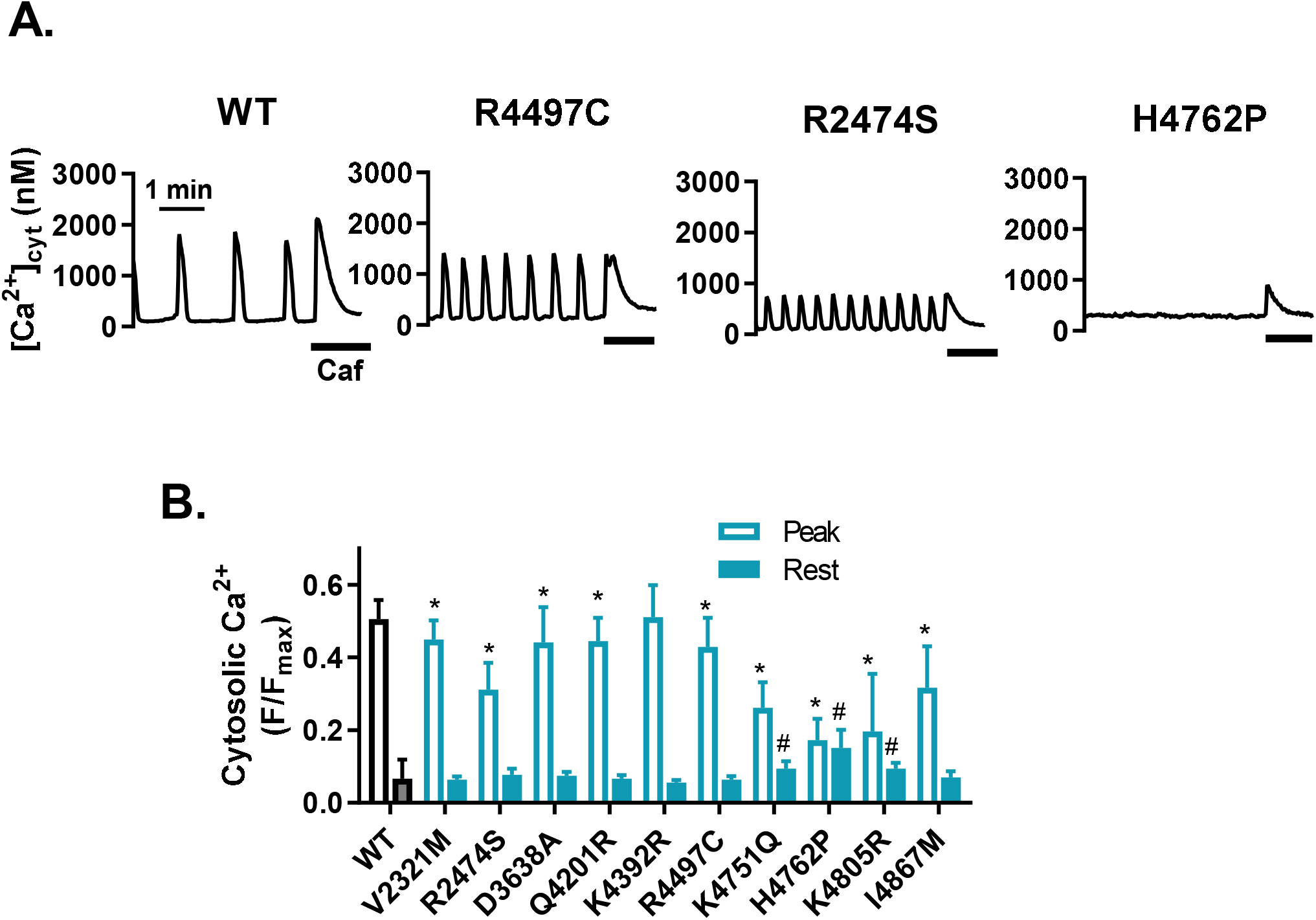
Calculated [Ca^2+^]_cyt_ in HEK293 cells expressing WT and mutant RyR2. **A.** Representative time course of [Ca^2+^]_cyt_ obtained by converting the fluo-4 signals in Fig. 2A into Ca^2+^ concentrations. Data were obtained in normal Krebs solution followed by application of 10 mM caffeine (black line). [Ca^2+^]_cy_ were obtained by converting from the data in Fig. 2A. **B**. Average peak and resting cytosolic [Ca^2+^]_cyt_. **C**. Relation between oscillation frequency and peak [Ca^2+^]_cyt_.

**Supplemental Figure 2.**
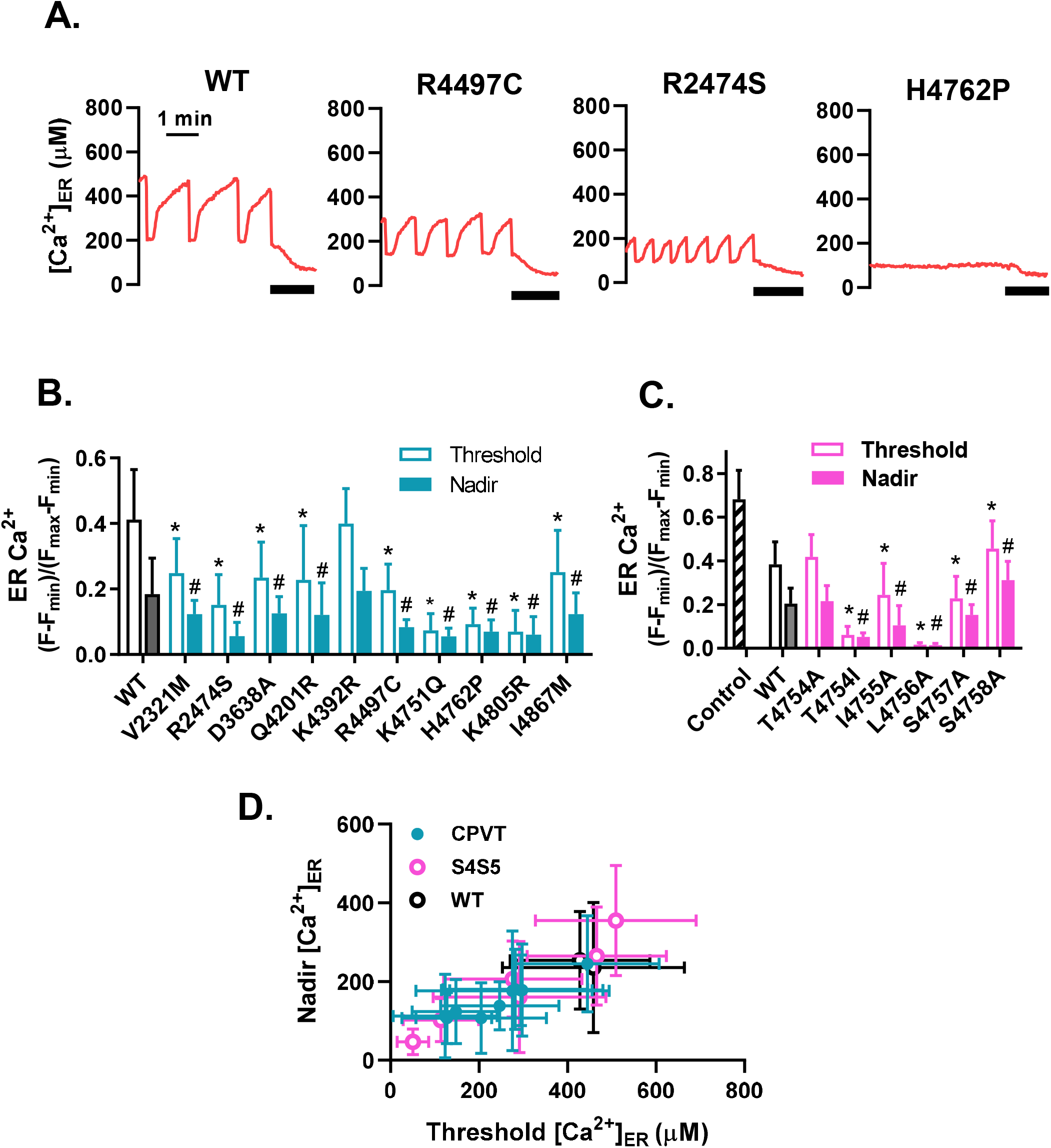
[Ca^2+^]_ER_ in HEK293 cells expressing WT and mutant RyR2. **A.** Time course of [Ca^2+^]_ER_ obtained by converting the R-CEPIA1_ER_ signals in Fig. 3A into Ca^2+^ concentrations. Data were obtained in normal Krebs solution followed by application of 10 mM caffeine (black line). **B**. Average threshold and nadir [Ca^2+^]_ER_ signals in cells expressing CPVT mutants. **C**. Average threshold and nadir [Ca^2+^]_ER_ signals in cells expressing artificial mutants. **D**. Relation between threshold and nadir [Ca^2+^]_ER_.

## References

Arakawa, J., A. Hamabe, T. Aiba, T. Nagai, M. Yoshida, T. Touya, N. Ishigami, H. Hisadome, S. Katsushika, H. Tabata, Y. Miyamoto, and W. Shimizu. 2015. A novel cardiac ryanodine receptor gene (RyR2) mutation in an athlete with aborted sudden cardiac death: a case of adult-onset catecholaminergic polymorphic ventricular tachycardia. Heart Vessels. 30:835–840.

Bassani, J.W., W. Yuan, and D.M. Bers. 1995. Fractional SR Ca release is regulated by trigger Ca and SR Ca content in cardiac myocytes. Am J Physiol. 268:C1313–1319.

Bers D.M. 2002. Cardiac excitation-contraction coupling. Nature. 415:198–205.

Bers, D.M., C.W. Patton, and R. Nuccitelli. 2010. A practical guide to the preparation of Ca^2+^ buffers. Methods Cell Biol. 99:1–26.

Bovo, E., J.L. Martin, J. Tyryfter, P.P. de Tombe, and A.V. Zima. 2016. R-CEPIA1er as a new tool to directly measure sarcoplasmic reticulum [Ca] in ventricular myocytes. Am J Physiol Heart Circ Physiol. 311:H268–275.

Cheung W.Y. 1980. Calmodulin plays a pivotal role in cellular regulation. Science. 207:19–27.

Chugun, A., O. Sato, H. Takeshima, and Y. Ogawa. 2007. Mg^2+^ activates the ryanodine receptor type 2 (RyR2) at intermediate Ca^2+^ concentrations. Am J Physiol Cell Physiol. 292:C535–544.

Endo, M. 1977. Calcium release from the sarcoplasmic reticulum. Physiol Rev. 57:71–108.

Fabiato, A. 1983. Calcium-induced release of calcium from the cardiac sarcoplasmic reticulum. Am J Physiol. 245:C1–14.

Fujii, Y., H. Itoh, S. Ohno, T. Murayama, N. Kurebayashi, H. Aoki, M. Blancard, Y. Nakagawa, S. Yamamoto, Y. Matsui, M. Ichikawa, K. Sonoda, T. Ozawa, K. Ohkubo, I. Watanabe, P. Guicheney, and M. Horie. 2017. A type 2 ryanodine receptor variant associated with reduced Ca^2+^ release and short-coupled torsades de pointes ventricular arrhythmia. Heart Rhythm. 14:98–107.

Gomez, A.C., and N. Yamaguchi. 2014. Two regions of the ryanodine receptor calcium channel are involved in Ca(2+)-dependent inactivation. Biochemistry. 53:1373–1379.

Guo, T., D. Gillespie, and M. Fill. 2012. Ryanodine receptor current amplitude controls Ca2+ sparks in cardiac muscle. Circ Res. 111:28–36.

Harkins, A.B., N. Kurebayashi, and S.M. Baylor. 1993. Resting myoplasmic free calcium in frog skeletal muscle fibers estimated with fluo-3. Biophys J. 65:865–881.

Jiang, D., R. Wang, B. Xiao, H. Kong, D.J. Hunt, P. Choi, L. Zhang, and S.R. Chen. 2005. Enhanced store overload-induced Ca^2+^ release and channel sensitivity to luminal Ca^2+^ activation are common defects of RyR2 mutations linked to ventricular tachycardia and sudden death. Circ Res. 97:1173–1181.

Jiang, D., B. Xiao, D. Yang, R. Wang, P. Cho, L. Zhang, H. Cheng, and S.R.W. Chen. 2004. RyR2 mutations linked to ventricular tachycardia and sudden death reduce the threshold for store-overload-induced Ca^2+^ release (SOICR). Proc Natl Acad Sci U S A. 101:13062–13067.

Jiang, D., B. Xiao, L. Zhang, and S.W. Chen. 2002. Enhanced Basal Activity of a Cardiac Ca^2+^ Release Channel (Ryanodine Receptor) Mutant Associated With Ventricular Tachycardia and Sudden Death. Circulation Research. 91:218–225.

Jones, P.P., D. Jiang, J. Bolstad, D.J. Hunt, L. Zhang, N. Demaurex, and S.R. Chen. 2008. Endoplasmic reticulum Ca^2+^ measurements reveal that the cardiac ryanodine receptor mutations linked to cardiac arrhythmia and sudden death alter the threshold for store-overload-induced Ca^2+^ release. Biochem J. 412:171–178.

Kawamura, M., S. Ohno, N. Naiki, I. Nagaoka, K. Dochi, Q. Wang, K. Hasegawa, H. Kimura, A. Miyamoto, Y. Mizusawa, H. Itoh, T. Makiyama, N. Sumitomo, H. Ushinohama, K. Oyama, N. Murakoshi, K. Aonuma, H. Horigome, T. Honda, M. Yoshinaga, M. Ito, and M. Horie. 2013. Genetic Background of Catecholaminergic Polymorphic Ventricular Tachycardia in Japan. Circulation Journal. 77:1705–1713.

Laitinen, P.J., K.M. Brown, K. Piippo, H. Swan, J.M. Devaney, B. Brahmbhatt, E.A. Donarum, M. Marino, N. Tiso, M. Viitasalo, L. Toivonen, D.A. Stephan, and K. Kontula. 2001. Mutations of the cardiac ryanodine receptor (RyR2) gene in familial polymorphic ventricular tachycardia. Circulation. 103:485–490.

Lakatta E.G. 1992. Functional implications of spontaneous sarcoplasmic reticulum Ca^2+^ release in the heart. Cardiovasc Res. 26:193–214.

Lieve, K.V.V., J.M.A. Verhagen, J. Wei, J.M. Bos, C. van der Werf, I.N.F. Roses, G.M.S. Mancini, W. Guo, R. Wang, F. van den Heuvel, I.M.E. Frohn-Mulder, W. Shimizu, A. Nogami, H. Horigome, J.D. Roberts, A. Leenhardt, H.J.G. Crijns, A.C. Blank, T. Aiba, A.C.P. Wiesfeld, N.A. Blom, N. Sumitomo, J. Till, M.J. Ackerman, S.R.W. Chen, I. van de Laar, and A.A.M. Wilde. 2019. Linking the heart and the brain: Neurodevelopmental disorders in patients with catecholaminergic polymorphic ventricular tachycardia. Heart Rhythm. 16:220–228.

Lukyanenko, V., I. Gyorke, and S. Gyorke. 1996. Regulation of calcium release by calcium inside the sarcoplasmic reticulum in ventricular myocytes. Pflugers Arch. 432:1047–1054.

Means, S., A.J. Smith, J. Shepherd, J. Shadid, J. Fowler, R.J. Wojcikiewicz, T. Mazel, G.D. Smith, and B.S. Wilson. 2006. Reaction diffusion modeling of calcium dynamics with realistic ER geometry. Biophys J. 91:537–557.

Medeiros-Domingo, A., Z.A. Bhuiyan, D.J. Tester, N. Hofman, H. Bikker, J.P. van Tintelen, M.M. Mannens, A.A. Wilde, and M.J. Ackerman. 2009. The RYR2-encoded ryanodine receptor/calcium release channel in patients diagnosed previously with either catecholaminergic polymorphic ventricular tachycardia or genotype negative, exercise-induced long QT syndrome: a comprehensive open reading frame mutational analysis. J Am Coll Cardiol. 54:2065–2074.

Murayama, T., and N. Kurebayashi. 2011. Two ryanodine receptor isoforms in nonmammalian vertebrate skeletal muscle: possible roles in excitation-contraction coupling and other processes. Prog Biophys Mol Biol. 105:134–144.

Murayama, T., N. Kurebayashi, M. Ishigami-Yuasa, S. Mori, Y. Suzuki, R. Akima, H. Ogawa, J. Suzuki, K. Kanemaru, H. Oyamada, Y. Kiuchi, M. Iino, H. Kagechika, and T. Sakurai. 2018. Efficient High-Throughput Screening by Endoplasmic Reticulum Ca^2+^ Measurement to Identify Inhibitors of Ryanodine Receptor Ca^2+^-Release Channels. Mol Pharmacol. 94:722–730.

Murayama, T., N. Kurebayashi, T. Oba, H. Oyamada, K. Oguchi, T. Sakurai, and Y. Ogawa. 2011. Role of amino-terminal half of the S4-S5 linker in type 1 ryanodine receptor (RyR1) channel gating. J Biol Chem. 286:35571–35577.

Murayama, T., N. Kurebayashi, H. Ogawa, T. Yamazawa, H. Oyamada, J. Suzuki, K. Kanemaru, K. Oguchi, M. Iino, and T. Sakurai. 2016. Genotype-Phenotype Correlations of Malignant Hyperthermia and Central Core Disease Mutations in the Central Region of the RYR1 Channel. Hum Mutat. 37:1231–1241.

Murayama, T., N. Kurebayashi, and Y. Ogawa. 2000. Role of Mg^2+^ in Ca^2+^-induced Ca^2+^ release through ryanodine receptors of frog skeletal muscle: modulations by adenine nucleotides and caffeine. Biophys J. 78:1810–1824.

Murayama, T., N. Kurebayashi, T. Yamazawa, H. Oyamada, J. Suzuki, K. Kanemaru, K. Oguchi, M. Iino, and T. Sakurai. 2015. Divergent Activity Profiles of Type 1 Ryanodine Receptor Channels Carrying Malignant Hyperthermia and Central Core Disease Mutations in the Amino-Terminal Region. PLoS One. 10:e0130606.

Nelson, F.E., S. Hollingworth, L.C. Rome, and S.M. Baylor. 2014. Intracellular calcium movements during relaxation and recovery of superfast muscle fibers of the toadfish swimbladder. J Gen Physiol. 143:605–620.

Nishio, H., M. Iwata, A. Tamura, T. Miyazaki, K. Tsuboi, and K. Suzuki. 2008. Identification of a novel mutation V2321M of the cardiac ryanodine receptor gene of sudden unexplained death and a phenotypic study of the gene mutations. Leg Med (Tokyo). 10:196–200.

Nozaki, Y., Y. Kato, K. Uike, K. Yamamura, M. Kikuchi, M. Yasuda, S. Ohno, M. Horie, T. Murayama, N. Kurebayashi, and H. Horigome. 2020. Co-Phenotype of Left Ventricular Non-Compaction Cardiomyopathy and Atypical Catecholaminergic Polymorphic Ventricular Tachycardia in Association With R169Q, a Ryanodine Receptor Type 2 Missense Mutation. Circ J. 84:226–234.

Postma, A.V., I. Denjoy, J. Kamblock, M. Alders, J.M. Lupoglazoff, G. Vaksmann, L. Dubosq-Bidot, P. Sebillon, M.M. Mannens, P. Guicheney, and A.A. Wilde. 2005. Catecholaminergic polymorphic ventricular tachycardia: RYR2 mutations, bradycardia, and follow up of the patients. J Med Genet. 42:863–870.

Priori, S.G., and S.R. Chen. 2011. Inherited dysfunction of sarcoplasmic reticulum Ca2+ handling and arrhythmogenesis. Circ Res. 108:871–883.

Priori, S.G., C. Napolitano, M. Memmi, B. Colombi, F. Drago, M. Gasparini, L. DeSimone, F. Coltorti, R. Bloise, R. Keegan, F.E. Cruz Filho, G. Vignati, A. Benatar, and A. DeLogu. 2002. Clinical and molecular characterization of patients with catecholaminergic polymorphic ventricular tachycardia. Circulation. 106:69–74.

Priori, S.G., C. Napolitano, N. Tiso, M. Memmi, G. Vignati, R. Bloise, V. Sorrentino, and G.A. Danieli. 2001. Mutations in the cardiac ryanodine receptor gene (hRyR2) underlie catecholaminergic polymorphic ventricular tachycardia. Circulation. 103:196–200.

Qin, J., G. Valle, A. Nani, A. Nori, N. Rizzi, S.G. Priori, P. Volpe, and M. Fill. 2008. Luminal Ca2+ regulation of single cardiac ryanodine receptors: insights provided by calsequestrin and its mutants. J Gen Physiol. 131:325–334.

Rios, E. 2018. Calcium-induced release of calcium in muscle: 50 years of work and the emerging consensus. J Gen Physiol. 150:521–537.

Robertson, S.P., J.D. Johnson, and J.D. Potter. 1981. The time-course of Ca2+ exchange with calmodulin, troponin, parvalbumin, and myosin in response to transient increases in Ca2+. Biophys J. 34:559–569.

Satoh, H., L.A. Blatter, and D.M. Bers. 1997. Effects of [Ca2+]i, SR Ca2+ load, and rest on Ca2+ spark frequency in ventricular myocytes. Am J Physiol. 272:H657–668.

Shannon, T.R., F. Wang, J. Puglisi, C. Weber, and D.M. Bers. 2004. A mathematical treatment of integrated Ca dynamics within the ventricular myocyte. Biophys J. 87:3351–3371.

Sitsapesan, R., and A.J. Williams. 1997. Regulation of current flow through ryanodine receptors by luminal Ca2+. J Membr Biol. 159:179–185.

Suzuki, J., K. Kanemaru, K. Ishii, M. Ohkura, Y. Okubo, and M. Iino. 2014. Imaging intraorganellar Ca^2+^ at subcellular resolution using CEPIA. Nat Commun. 5:4153.

Tester, D.J., D.B. Spoon, H.H. Valdivia, J.C. Makielski, and M.J. Ackerman. 2004. Targeted mutational analysis of the RyR2-encoded cardiac ryanodine receptor in sudden unexplained death: a molecular autopsy of 49 medical examiner/coroner’s cases. Mayo Clin Proc. 79:1380–1384.

Tsien, R.W., R.S. Kass, and R. Weingart. 1979. Cellular and subcellular mechanisms of cardiac pacemaker oscillations. J Exp Biol. 81:205–215.

Uehara, A., T. Murayama, M. Yasukochi, M. Fill, M. Horie, T. Okamoto, Y. Matsuura, K. Uehara, T. Fujimoto, T. Sakurai, and N. Kurebayashi. 2017. Extensive Ca^2+^ leak through K4750Q cardiac ryanodine receptors caused by cytosolic and luminal Ca^2+^ hypersensitivity. J Gen Physiol. 149:199–218.

Wehrens, X.H., S.E. Lehnart, F. Huang, J.A. Vest, S.R. Reiken, P.J. Mohler, J. Sun, S. Guatimosim, L.S. Song, N. Rosemblit, J.M. D’Armiento, C. Napolitano, M. Memmi, S.G. Priori, W.J. Lederer, and A.R. Marks. 2003. FKBP12.6 deficiency and defective calcium release channel (ryanodine receptor) function linked to exercise-induced sudden cardiac death. Cell. 113:829–840.

Xiao, Z., W. Guo, B. Sun, D.J. Hunt, J. Wei, Y. Liu, Y. Wang, R. Wang, P.P. Jones, T.G. Back, and S.R. Chen. 2016. Enhanced Cytosolic Ca^2+^ Activation Underlies a Common Defect of Central Domain Cardiac Ryanodine Receptor Mutations Linked to Arrhythmias. J Biol Chem. 291:24528–24537.

Zhao, Y., S. Araki, J. Wu, T. Teramoto, Y.F. Chang, M. Nakano, A.S. Abdelfattah, M. Fujiwara, T. Ishihara, T. Nagai, and R.E. Campbell. 2011. An expanded palette of genetically encoded Ca^2+^ indicators. Science. 333:1888–1891.

